# Evidence of RNA polymerase III recruitment and transcription at protein-coding gene promoters

**DOI:** 10.1101/2024.06.08.598009

**Authors:** K C Rajendra, Ruiying Cheng, Sihang Zhou, Simon Lizarazo, Duncan Smith, Kevin Van Bortle

## Abstract

RNA polymerase (Pol) I, II, and III are most commonly described as having distinct roles in synthesizing ribosomal RNA (rRNA), messenger RNA (mRNA), and specific small noncoding (nc)RNAs, respectively. This delineation of transcriptional responsibilities is not definitive, however, as evidenced by instances of Pol II recruitment to genes conventionally transcribed by Pol III, including the co-transcription of *RPPH1* - the catalytic RNA component of RNase P. A comprehensive understanding of the interplay between RNA polymerase complexes remains lacking, however, due to limited comparative analyses for all three enzymes. To address this gap, we applied a uniform framework for quantifying global Pol I, II, and III occupancies that integrates currently available human RNA polymerase ChIP-seq datasets. Occupancy maps are combined with a comprehensive multi-class promoter set that includes protein-coding genes, noncoding genes, and repetitive elements. While our genomic survey appropriately identifies recruitment of Pol I, II, and III to canonical target genes, we unexpectedly discover widespread recruitment of the Pol III machinery to promoters of specific protein-coding genes, supported by colocalization patterns observed for several Pol III-specific subunits. We show that Pol III-occupied Pol II promoters are enriched for small, nascent RNA reads terminating in a run of 4 Ts, a unique hallmark of Pol III transcription termination and evidence of active Pol III activity at these sites. Pol III disruption differentially modulates the expression of Pol III-occupied coding genes, which are functionally enriched for ribosomal proteins and genes broadly linked to unfavorable outcomes in cancer. Our map also identifies additional, currently unannotated genomic elements occupied by Pol III with clear signatures of nascent RNA species that are sensitive to disruption of La (SSB) - a Pol III-related RNA chaperone protein. These findings reshape our current understanding of the interplay between Pols II and III and identify potentially novel small ncRNAs with broad implications for gene regulatory paradigms and RNA biology.

## Introduction

RNA polymerase (Pol) I, II, and III are recruited by distinct transcription factor (TF) repertoires that functionally sub-divide the recruitment and ensuing transcription potential of each enzyme. However, the distinctions between these TF repertoires and the transcriptional machineries themselves are blurred by the fact that multiple subunits and individual regulatory factors are shared by two or more polymerases^1–3^. Taken further, the simple paradigm of transcriptional division itself is challenged by examples of Pol II overlap reported at both Pol I and Pol III-transcribed genes^4–6^, including examples of shared transcription^7^. The full extent of RNA polymerase overlap and shared transcription remains largely unexplored, nevertheless, due to limited genomic studies of Pol I and Pol III in humans.

Pol III is unique from other polymerases in multiple respects, including that it exclusively transcribes genes encoding small noncoding RNA (ncRNA), many of which are critical for cell growth and proliferation^8–10^. Examples include tRNA and 5S ribosomal RNA (rRNA) - both integral components of translation - as well as H1 RNA, the catalytic RNA component of RNase P^7^. The gene encoding H1, *RPPH1*, is transcribed by both Pol III and Pol II, exemplifying scenarios of competent recruitment and transcription by multiple RNA polymerase machineries^11^.

Recently, we explored the global occupancy of human Pol III by integrating ChIP-seq experiments for a large subset of Pol III subunits^12^. Defining the binding patterns of many Pol III subunits increases confidence of complex, rather than subunit-specific, chromatin signatures. We note that these signatures unexpectedly included numerous overlapping coordinates beyond the canonical annotation of the Pol III “transcriptome”, defined on the basis of previous biochemical and genomic mapping experiments^12^.

Here, we reexamine the canonical and noncanonical binding preferences of human Pol III by integrating multiple genomic approaches with comprehensive coding and noncoding gene annotations. We apply a uniform, context-agnostic framework for scoring Pol I, II, and III occupancies by integrating >200 ChIP-seq datasets, facilitating unbiased annotation of polymerase occupancy and overlap. While this approach recovers canonical recruitment and transcription patterns, we unexpectedly observe high Pol III scores at specific protein-coding gene promoters. These results inspired deeper consideration of the transcriptional competency and consequences of Pol III occupancy at these sites.

Pol III transcription termination relies on complementary binding between nascent RNA and a relatively short repeat of 4 to 6 Thymidine (T) sequences on the non-template DNA strand^13^. Thereafter, the terminal oligo(U) tract in the corresponding nascent RNA is recognized by SSB (La), an RNA chaperone protein important for Pol III-derived RNA stability and processing^14^. The Pol III termination signature is unique from Pol I, which relies on a relatively long T-tract^15^, as well as Pol II termination signals following cleavage and polyadenylation of mRNA^16^. We therefore developed a complementary framework for scoring global patterns of single nucleotide 3’-end pileup at minimal sequences of 4 Ts (referred to here as “T4”) as a molecular breadcrumb of Pol III transcription. These data, when coupled with Pol III occupancy patterns, provide compelling evidence of active Pol III transcription at a multitude of protein-coding gene promoters. Moreover, the Pol II-derived mRNA products from genes bound by Pol III are notably sensitive to Pol III disruption, altogether establishing a novel link between Pol III recruitment, transcription, and important downstream consequences on protein-coding genes.

## Results

### Global interrogation of RNA polymerase I, II, III occupancy

We first retrieved available ChIP-seq data corresponding to complex-specific Pol I, II, and III subunits. Previous mapping experiments for human Pol I are exceptionally limited, with only 2 datasets reported for binding of the large sub-units, POLR1A and POLR1B^17,18^. Pol II datasets are significantly more abundant, with 153 POLR2A ChIP-seq experiments available for the large subunit of Pol II. ChIP-seq experiments for Pol III, on the other hand, include a multitude of Pol III-specific subunits, including POLR3A, - B, -C, -D, -E, and -G (Figure 1a, Supplemental Table 1). The broader profile of Pol III subunit occupancy is meaningful for more confidently establishing *de facto* recruitment of the Pol III machinery.

**Figure 1.**
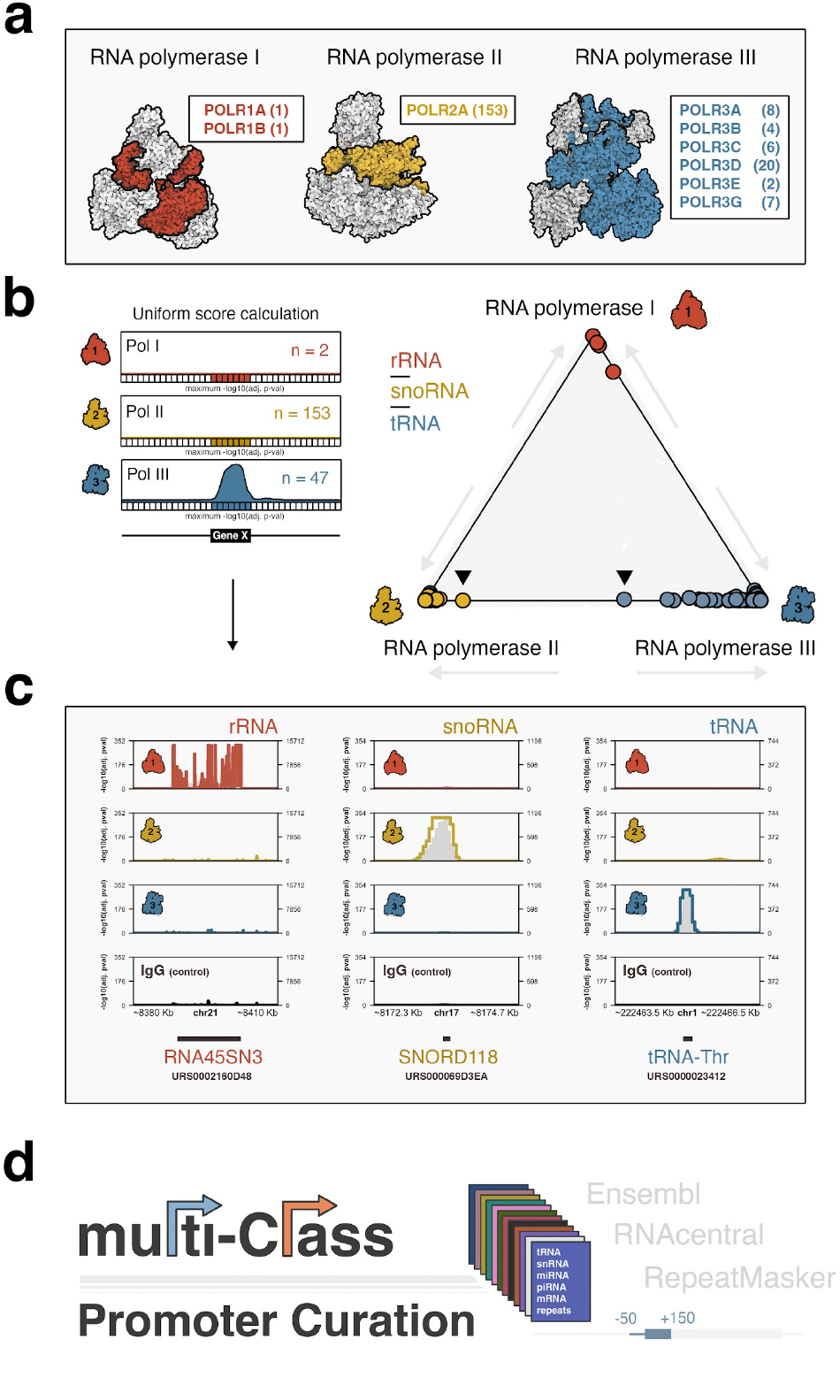
Genome-wide survey of human RNA polymerase binding patterns. **(a)** Illustrative overview of Pol I, II, and III complexes, coloring and labels correspond to individual subunits mapped by ChIP-seq. **(b)** Application of a uniform scoring framework (left) for composite Pol I, II, and III ChIP-seq signals. Pol occupancy scores are defined globally across 50 bp bins by assigning adjusted Poisson probabilities determined against maximum, bin-specific lambda scores. Global Pol scoring recovers canonical polymerase-gene dominance at rRNA (Pol I), snoRNA (Pol II), and tRNA (Pol III). **(c)** Specific examples of Pol I dominance at ribosomal RNA genes (RNA45SN3, left), Pol II dominance at snoRNA genes (SNORD118, center), and Pol III dominance at tRNA genes (tRNA-Thr, right). **(d)** Coding, noncoding, and repeat annotations were integrated into a multi-class promoter set, defined as a window 50 bp upstream to 150 downstream of transcription start sites, for global scoring of polymerase occupancies.

Pol II and III ChIP-seq data were compiled by unique cellular context for each individual subunit and uniformly scaled to the minimum number of subunit-specific sample reads. Summary profiles were then generated by combining all processed data and subsequently scaling to 100 million total reads, thereby producing a profile weighed equally by each biological context. Finally, a median Pol III summary profile was calculated across all individual subunits and rescaled to 100 million reads. Due to the limited experimental data for Pol I, POLR1A and POLR1B datasets were simply aggregated and analogously scaled to 100 million reads.

We next applied a peak calling framework for establishing global polymerase-specific scores at 50 bp resolution. Briefly, a Poisson probability was calculated for every 50 bp bin by comparing the observed signal to a local expectation - lambda - defined as the maximum polymerase signal at 5 Kb, 10 Kb, or genome-wide, a stringent approach that accounts for local chromatin biases specific to each genomic window^19^. We then compared Pol I, II, and III scores at canonical gene annotations, including rRNA, snoRNA, and tRNA. As expected, our uniform scoring framework successfully recovers Pol I-specific occupancy of rRNA, Pol II-specific occupancy of snoRNA, and Pol III-specific occupancy of tRNA (Figure 1b, specific examples shown in Figure 1c).

To effectively visualize variable levels of polymerase occupancy at these and other gene categories, we applied a center-of-mass equation to define the coordinate position of each specific gene, thereby establishing a tool for simultaneous three-way comparison of Pol I, II, and III “dominance”. Genes with strong signal for only a single polymerase will be positioned close to its respective enzyme (e.g., tRNA-Pol III, Figure 1b), whereas genes with strong signal for two machineries will be positioned closer to the center of a given axis. Using this approach, specific Pol III-transcribed tRNA genes include notable Pol II signal, consistent with previous reports, yet specific Pol II-transcribed snoRNA genes are also pulled, to some degree, from Pol II dominance towards Pol III (e.g., Figure 1b, arrows). We therefore expanded our survey and more closely examined significant Pol III binding events at both canonical and noncanonical genes.

To systematically annotate Pol I, II, and III patterns while also addressing the underlying challenges of variable gene annotations sets, we integrated coding and noncoding genes by combining Ensembl transcript annotations (GENCODE V43) with the RNAcentral noncoding RNA sequence data-base (v22.0). Gene promoter windows were uniformly defined as a 200 bp window, including 50 bp upstream and 150 bp downstream of each individual transcription start site, harmonized across instances of varied annotation. Additionally, intervals corresponding to repeat sequences annotated by RepeatMasker were incorporated into our curated, multi-class promoter resource (Figure 1d). We thereafter assigned the maximum score for each polymerase over a given genomic interval and identified protein-coding, noncoding, and repeat intervals with significant polymerase occupancy (adjusted p-value < 0.01).

### Pol III transcription and termination at canonical genes

Beyond tRNA and 5S rRNA, the Pol III transcriptome includes U6 and U6atac spliceosomal RNA, 7SK small nuclear (sn)RNA, 7SL signal recognition particle (SRP) RNA and its evolutionary derivatives - Alu, BC200, and snaR - as well as Y RNA, RNase P and RNase MRP RNA (RMRP, H1), Vault RNA, and nc866^8,10,20^. Recruitment of Pol III to each gene type is directed through either gene-internal promoter elements (e.g. 5S rRNA, tRNA) or upstream regulatory sequences that are recognized by specific transcription factor repertoires^21^. Consistent with this established annotation of the Pol III-transcriptome, we observe strong Pol III occupancy scores across most gene subclasses for Pol III (Figure 2a). However, whereas Pol III is the dominant polymerase at nearly all canonical genes, our framework recovers shared dominance of *RPPH1* by Pol II and Pol III as previously described (Supplemental Figure 1)^7^.

**Figure 2.**
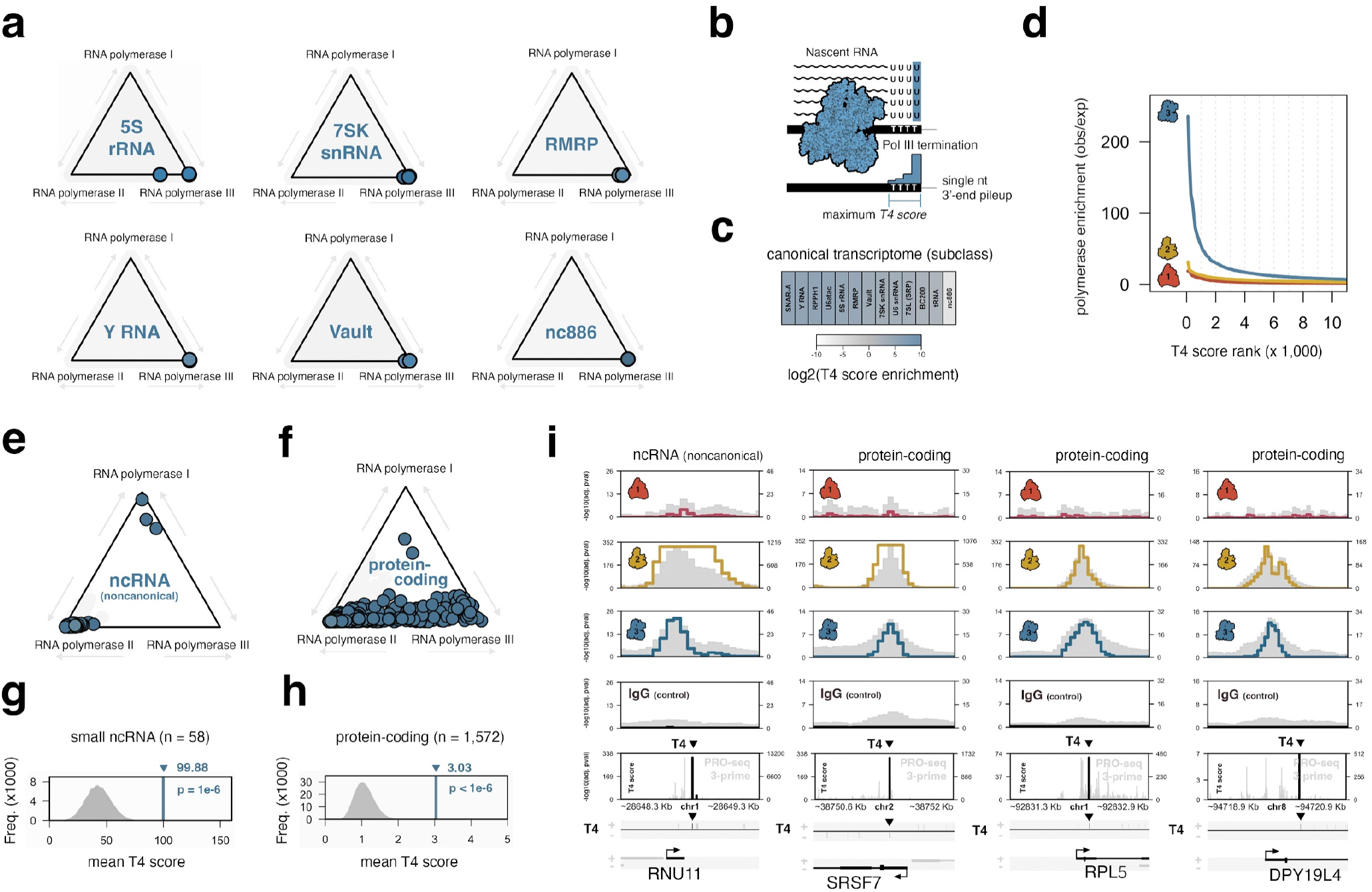
Discovery of RNA polymerase III occupancy and transcription termination signatures at noncanonical loci, including protein-coding gene promoters. **(a)** Analysis of polymerase signal dominance at specific subclasses of Pol III-transcribed genes, including 5S rRNA, 7SK snRNA, RMRP, Y RNA, Vault, and nc886. **(b)** Illustrative overview of Pol III termination, which occurs at a run of 4-6 Thymidine (T) sequences. T4 scoring of 3’-single nucleotide pileup in nascent RNA is used as a molecular readout of Pol III transcription termination. **(c)** T4 score enrichment for specific Pol III-transcribed gene subclasses, with respect to all sites with significant Pol III occupancy (padj < .05). **(d)** Pol I, II, and III signal enrichment at gene promoters, ranked by maximum T4 scores present within 350 bp of a given transcription start site. **(e)** RNA polymerase dominance plot for small noncoding RNA genes not previously defined as Pol III-transcribed but characterized by significant Pol III occupancy in our metamap. **(f)** Analogous dominance plot for protein-coding genes with significant Pol III occupancy. **(g-h)** Observations and corresponding empirical null distributions for T4 scores at noncanonical small RNA genes (related to Figure 2e) and protein-coding gene promoters (related to Figure 2f). **(i)** Visualization of Pol I, II, IIII occupancies at RNU11 - a small ncRNA gene canonically expressed by Pol II - and multiple protein-coding genes, including SRSF7, RPL5, and DPY19L4. All genes are defined as having significant Pol III occupancy and T4 score enrichment. Significant T4 score peaks are highlighted.

To further query Pol III activity signatures at canonical genes, we generated a complementary composite profile of precision nuclear run-on RNA-sequencing (PRO-seq), a method that isolates nascent RNA to measure transcription rather than steady-state RNA levels (Supplemental Table 2)^22^. Based on the unique termination mechanism of Pol III at 4-6 Ts, we considered that single-nucleotide read enrichment mapping to the terminal 3’ nucleotide of nascent RNA over 4 or more contiguous Ts would be a strong indicator of Pol III transcription termination (Figure 2b). We therefore applied a similar peak scoring framework to 3’-read pileup instances at T4 sequences, deriving a local lambda from the maximum signal intensity at either 20 or 50 bp surrounding each T4. Gene-specific T4 scores were then calculated by retrieving the maximum gene-internal T4 score within 350 bp of a transcription start site. Using this approach, we find that most Pol III-transcribed genes are significantly enriched for strong T4 signatures (Figure 2c). Interestingly, T4 enrichment patterns are notably lower for tRNA genes compared to other Pol III-transcribed gene classes, likely driven by rapid cleavage events of nascent tRNA 3’ trailer sequences^23^. Globally, we find that Pol III is highly enriched at genes ranked by maximum T4 score, in contrast to Pol I and Pol II (Figure 2d). These data indicate a clear association of nascent T4 enrichment patterns and Pol III activity, both at canonical target genes and potentially unexpected loci. We therefore turned our attention to the broader patterns of Pol III occupancy and transcription towards better understanding the global Pol III landscape.

### Pol III occupancy and transcription signatures at noncanonical small RNA and protein-coding genes

We first re-visited our RNA polymerase scoring framework, specifically searching for sites with significant Pol III levels beyond the canonical Pol III transcriptome. For all protein-coding transcription start sites, genomic intervals within 500 bp of an established Pol III-transcribed gene or a similar small ncRNA subclass were masked to avoid potential distance-related artifacts. Altogether, this approach identifies 58 noncoding gene promoters and, surprisingly, 1,572 protein-coding gene promoters with significant Pol III signal. Whereas most noncanonical ncRNA genes are dominated by Pol II (Figure 2e), we unexpectedly observe a continuum of Pol II-Pol III dominance at protein-coding gene promoters, with examples of Pol III dominance over Pol II (Figure 2f).

By surveying the T4 termination signatures at non-canonical Pol III loci, we further uncover significant T4 score enrichment within Pol III-occupied small ncRNA and protein-coding genes, strongly indicative of Pol III transcription at these loci (Figure 2g-h). Accordantly, single-nucleotide T4 pileup scores are enriched in Pol III-occupied genes, most significantly within the first ∼ 100 bp, suggesting that Pol III activity is restricted to a short genomic interval, likely constrained by the frequency of T4 sequence events (Supplemental Figure 2). Visualization of individual non-canonical loci, such as *RNU11* - a Pol II-derived snRNA component of the minor spliceosome - and protein-coding genes such as *SRSF7, RPL5*, and *DPY19L4*, illustrates this general trend of Pol III occupancy and sharp T4 pileups a short distance from transcription start sites (Figure 2i).

In addition to T4 score enrichment, we considered whether specific transcription factors (TFs) were enriched at Pol III-occupied protein-coding genes. TF-binding patterns were retrieved from ChIP-atlas^24^ and corresponding enrichment patterns determined by comparison against a randomization-based null model. Examination of TFs with established roles in Pol III recruitment reveals notable enrichment for BRF1, a TFIIIB subcomponent involved in recruiting Pol III to type 1 and type 2 promoters^21^, and SNAPC4, a SNAP-complex protein involved in recruiting Pol III to type 3 promoters (Supplemental Figure 3)^21^. We therefore similarly applied our uniform framework for scoring composite ChIP-seq profiles to Pol III-related transcription factors, including TFIIIB subunits (BRF1, TBP), SNAPC subunits (SNAPC1, -2, -4, -5), and TFIIIC subunits (GTF3C-1, -2, -5). Consistent with the overlap enrichment results, this method uncovers a strong correlation between Pol III and BRF1 scores at protein-coding gene promoters, with comparatively lower correlation scores observed for TBP, SNAPC4, and GTF3C5, and minimal scores for other TFIIIC and SNAPC subunits (Supplemental Figure 3). Examination of individual gene promoters further supports these trends, with evidence of BRF1 occupancy at specific protein-coding genes, such as *RPS7* (ribosomal protein S7), which features significant T4 scores indicative of Pol III termination.

**Figure 3.**
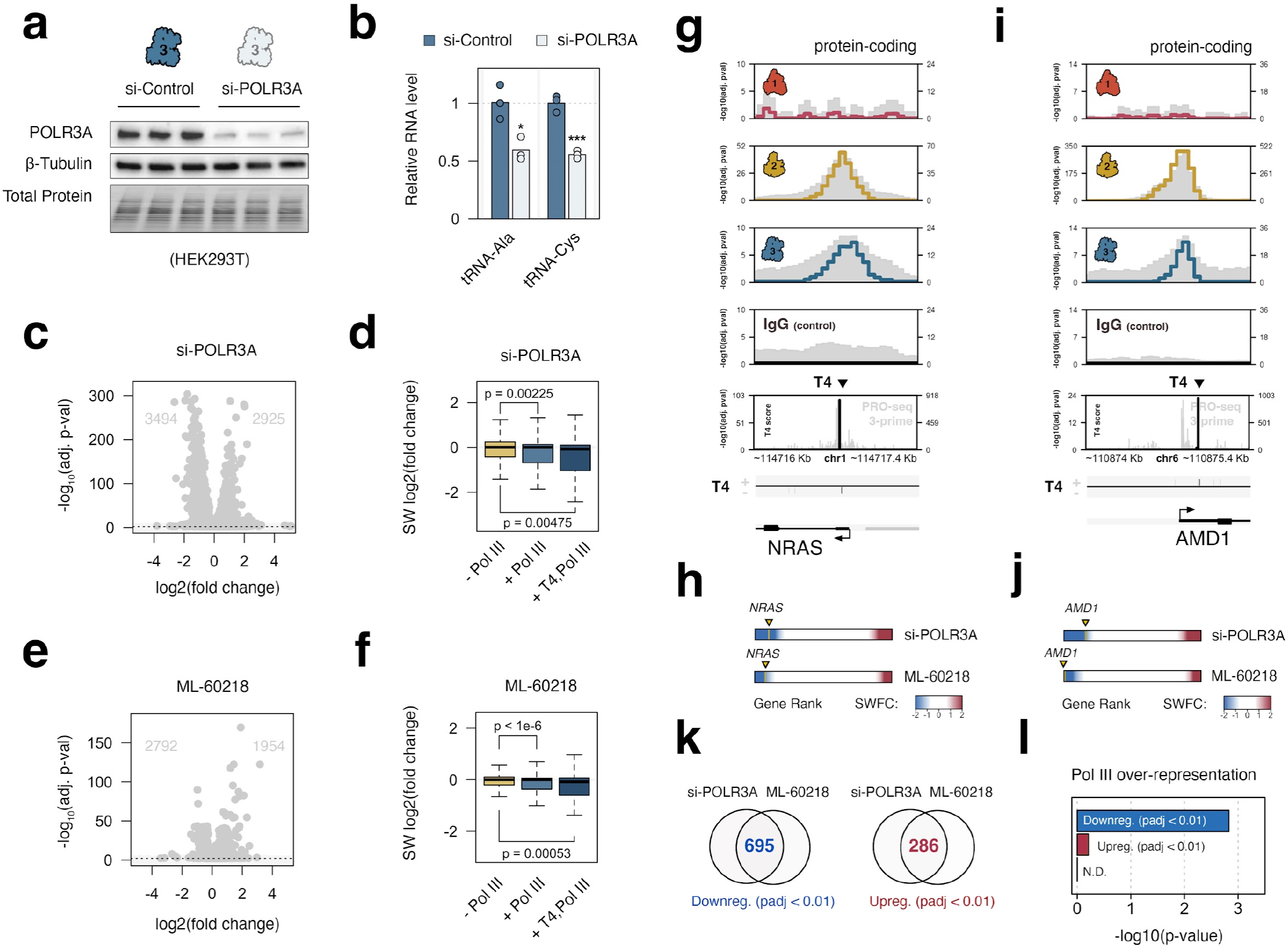
Pol III-occupied protein-coding gene promoters are sensitive to Pol III depletion and inhibition. **(a)** Immunoblot analysis POLR3A (RPC1) and beta-Tubulin protein levels in control (scramble) and POLR3A siRNA conditions in HEK293T cells. **(b)** RT-qPCR analysis of the relative change in Pol III-transcribed tRNA levels following siRNA-mediated reduction in HEK293T. **(c)** Differential expression profile for protein coding genes, including the number of up- and down-regulated genes (FDR < 0.05 and abs(Log2FC > 1) following disruption of POLR3A. **(d)** Distributions of significance-weighted fold change scores for protein-coding genes without Pol III, with Pol III, or with both Pol III and significant T4 scores. **(e)** Differential expression profile for protein coding genes, including the number of up- and down-regulated genes (FDR < 0.05 and abs(Log2FC > 1) following Pol III inhibition with small molecule ML-60218. **(f)** Distributions of significance-weighted fold change scores for protein-coding genes without Pol III, with Pol III, or with both significant Pol III and T4 scores, following Pol III inhibition with small molecule ML-60218. **(g-h)** Specific example of shared Pol II and Pol III occupancy at a protein-coding gene *NRAS* **(g)** which is highly sensitive to Pol III disruption **(h)**, either through depletion of POLR3A or through inhibition of Pol III with ML-60218. **(i-j)** Additional example of Pol III occupancy **(i)** and and Pol III sensitivity **(j)** at protein-coding gene *AMD1*. **(k)** Total number of shared differentially down-(blue) and upregulated genes (red) following si-POLR3A or ML-60218 (padj < 0.01). **(l)** Overlap enrichment of Pol III occupancy at down-, up-, and nondifferential (N.D.) gene sets

### Pol III occupied protein-coding genes are differentially sensitive to Pol III depletion and inhibition

The discovery of Pol III occupancy and transcription termination signatures led us to investigate the effect of Pol III disruption on these and other protein-coding gene sub-populations. Knockdown of *POLR3A* - the gene encoding Pol III large subunit RPC1 - is sufficient to significantly reduce Pol III-derived gene products, including multiple tRNAs (Figure 3a-b). We show that the consequence of *POLR3A* disruption extends far beyond canonical Pol III products, resulting in upregulation and downregulation of several thousand protein-coding genes (Figure 3c). Analysis of mRNA dynamics corresponding to genes occupied by Pol III, or occupied by Pol III with evidence of transcription termination, shows that reduction of Pol III generally reduces, rather than increases, the expression of protein-coding genes occupied by Pol III compared with genes lacking Pol III occupancy (Figure 3d).

As a complementary approach, we disrupted Pol III function with ML-60218 - a small molecule Pol III-specific inhibitor that we show drives upregulation and down-regulation of thousands of genes within an acute, 4-hour exposure (Figure 3e)^25^. As a distribution, Pol III-occupied protein-coding genes generally decrease in response to ML-60218, consistent with the biased effect of *POLR3A* depletion (Figure 3f).

In support of this trend, examination of specific Pol III-occupied genes illustrates examples of Pol III-sensitive protein-coding genes. For example, *NRAS* - a proto-oncogene encoding a Ras-family GTPase implicated in the progression of several forms of cancer^26^ - is included among the list of si-POLR3A and ML-60218 sensitive genes, with strong evidence of Pol III occupancy and signatures consistent with Pol III termination (Figure 3g-h). Similarly, *AMD1*, which encodes an adenosylmethionine decarboxylase implicated in multiple forms of cancer progression^27–29^, is characterized by strong evidence of Pol III occupancy and termination, and sensitive to si-POLR3A and ML-60218 exposure (Figure 3i-j).

We next considered whether significant Pol III-binding events, defined by our global occupancy map, were over-represented at Pol III-sensitive genes. We reasoned that genes determined to be significantly differential in both si-POLR3A and ML-60218 experiments are most likely to represent true, primary Pol III-sensitive features. Altogether, 695 genes are significantly downregulated in both conditions, and 286 significantly upregulated (Figure 3k; adjusted p-value < 0.01 in both conditions). Integration of these Pol III-sensitive gene sets with Pol III occupancy signatures uncovers significant over-representation of Pol III-binding, specifically at genes downregulated in both si-POLR3A and ML-60218 experiments (Figure 3l). Although Pol III-occupied loci continue to possess higher T4 scores independent of a particular gene group, the enrichment of T4 scores at genes that are downregulated by si-POLR3A and ML-60218 does not reach statistical significance (Supplemental Figure 4). Nevertheless, the enrichment of Pol III occupancy at downregulated genes, which contrasts that of both up-regulated and nondifferential gene sets, further supports a model in which Pol III recruitment and transcription may functionally enhance Pol II-mediated production of full-length mRNAs from these loci.

**Figure 4.**
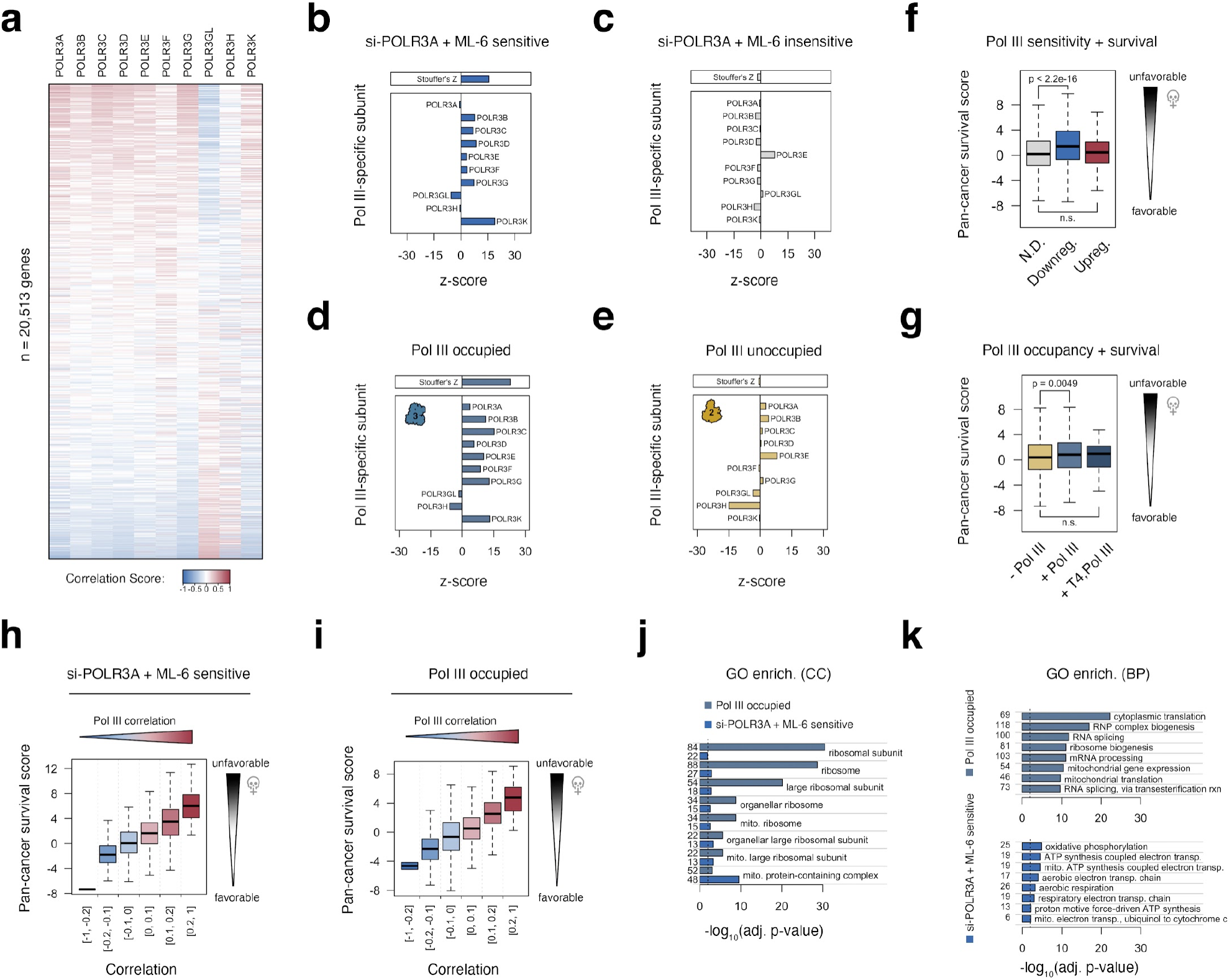
Expression of Pol III-sensitive and Pol III-occupied protein-coding genes is broadly linked with Pol III subunit expression and unfavorable outcomes in cancer. **(a)** Heatmap visualization of gene expression correlation scores for > 20,000 genes, ranked by the median correlation with Pol III-specific subunits. **(b-c)** Analysis of subunit-specific correlation features for genes downregulated in response to both si-POLR3A and ML-60218 experiments **(b)**, and genes that are non-differential in both experiments **(c)**. Barplots illustrate the strength and direction of correlation between a given gene set and genes encoding individual Pol III subunits, benchmarked against empirical null distributions. A summary z-score, calculated using Stouffer’s method, is represented at the top. **(d-e)** Analogous comparison of subunit-specific correlation features for Pol III occupied **(d)** and Pol III unoccupied **(e)** protein-coding genes. **(f)** Distributions of pan-cancer survival statistics related to the expression of genes that are nondifferential (N.D.), downregulated by both si-POLR3A and ML-60218 (downreg.), or upregulated in both conditions (upreg.). Survival statistics were retrieved from tcga-survival.com. **(g)** Distributions of pan-cancer survival statistics related to the expression of genes that Pol III unoccupied (-Pol III), Pol III occupied (+ Pol III), or Pol III occupied with T4 signatures (+ T4, Pol III). **(h-i)** Distributions of pan-cancer survival statistics related to the expression of genes as a function of its median correlation with Pol III-specific subunits (related to a) for Pol III-sensitive **(h)** or Pol III-occupied **(i)** genes. **(j)** Gene ontology (GO) enrichment analysis for cellular component (CC) terms significant (FDR < 0.05) in both Pol III-sensitive and Pol III-occupied gene sets. **(k)** Enriched biological process (BP) GO terms for Pol III-sensitive (top) and Pol III-occupied (bottom) gene sets.

### Pol III-sensitive and Pol III-occupied protein-coding genes are linked with Pol III correlation signatures across cancer

Building on our observations of Pol III occupancy and Pol III sensitivity at protein-coding genes, we further explored the relationship between Pol III and gene expression signatures captured by The Cancer Genome Atlas (TCGA) across thousands of diverse primary tumors and cancers^30^. We specifically examined gene expression correlates for Pol III-specific subunits *POLR3A, POLR3B, POLR3C, POLR3D, POLR3E, POLR3F, POLR3G, POLR3GL, POLR3H*, and *POLR3K* with more than 20,000 genes (Figure 4a). Though RNA levels are an imperfect representation of Pol III machinery availability in any particular context, the expansive breadth of expression signatures across thousands of samples offers a reasonably confident indication, collectively, of the variable Pol III levels for the purpose of linking specific genes to Pol III. Using this approach, we identify distinct subpopulations of protein-coding genes that are strongly associated, either positively or negatively, with the expression of most Pol III-specific subunits (Figure 4a).

To determine whether Pol III-sensitive and Pol III-occupied genes are significantly linked in the multi-cancer gene expression map, we compared the observed correlation scores for Pol III subunits individually using a randomization-based null model. Revisiting the 695 si-POLR3A and ML-60218 sensitive genes, we uncover a strong positive association with the expression of most Pol III-specific subunits, whereas si-POLR3A and ML-60218 insensitive genes are largely unrelated (Figure 4b-c). Similarly, we identify an even stronger positive relationship between Pol III subunit expression with protein-coding genes occupied by Pol III in our survey, in clear contrast to Pol III unoccupied loci (Figure 4d-e). These data provide further evidence that Pol III occupancy likely contributes to the expression landscape of specific protein-coding genes.

### Pol III-sensitive and Pol III-occupied protein-coding genes are functionally enriched for ribosomal proteins and genes linked to unfavorable outcomes in cancer

Pol III overactivity, which is ostensibly associated with its expansion to noncanonical loci such as protein-coding genes, has been described as a hallmark of cancer^31–33^. We therefore considered whether Pol III-sensitive, Pol III-occupied, and/or Pol III-correlated gene features are relevant to cancer outcomes in patients. We specifically considered the relationship between gene expression and clinical outcomes matched to specific primary tumors, as previously reported (tcga-survival.com^34^). Using this approach, we find that the distribution of genes downregulated by both si-POLR3A and ML-60218 have significantly higher z-scores, which reflect an indication of unfavorable outcomes among cancer patients, compared to si-POLR3A and ML-60218 insensitive genes (Figure 4f). Notably, this pattern is not observed for genes that are upregulated following Pol III disruption. To a lesser extent, we find that the distribution of Pol III-occupied protein-coding genes are also characterized as having significantly higher z-scores compared to Pol III-unoccupied genes (Figure 4g). However, further subdivideing Pol III-sensitive (downregulated) and Pol III-occupied protein-coding genes by their observed correlation patterns across cancer reveals a remarkable link between positively-correlated genes and negative patient outcomes (Figure 4h-i). These data give rise to a model in which Pol III recruitment and enhanced Pol II transcription of specific protein-coding genes may promote cellular growth and, consequently, disease progression.

To further explore whether specific gene functions are enriched in Pol III-related protein-coding genes, we performed gene ontology (GO) enrichment analysis on both Pol III-occupied and Pol III-sensitive gene sets. Both sets share significant enrichment for genes encoding ribosomal sub-units, with Pol III-occupied genes most strongly enriched for these proteins (Figure 4j). While this is further reflected in Pol III-occupied genes being enriched for “cytoplasmic translation” and “ribosome biogenesis”, Pol III-sensitive genes are instead primarily enriched for mitochondrial related metabolic processes, including “oxidative phosphorylation” and “ATP synthesis coupled electron transport” (Figure 4k). These results suggest that Pol III-sensitive and Pol III-occupied genes are closely related to biosynthesis and energy metabolism directly linked to cellular growth.

### Pol III recruitment and transcription at protein-coding genes does not produce stable small RNA species

Given the co-association of Pol III occupancy and T4 termination signatures at specific protein-coding genes, we naturally questioned whether Pol III activity at these sites produce stable small RNA species, hypothetically giving rise to a nested Pol III transcriptome derived from these regions. We therefore profiled small RNA abundance linked to Pol III-occupied protein-coding genes by integrating all human multi-tissue small RNA-seq data generated by the ENCODE project^35^ (Supplemental Table 3). Pol III-occupied protein-coding gene promoters are indeed characterized by significantly higher small RNA signatures, particularly for promoters with Pol III-binding and T4 signatures (Figure 5a). We find that this trend is further supported by small RNA-seq in HEK293T cells (Figure 5b). However, the distribution of small RNA abundance at these loci is minimal in comparison to canonical Pol III-transcribed genes (i.e. tRNA, Y RNA, vault RNA, etc.), suggesting that Pol III activity at these sites does not produce similarly stable ncRNA species (Figure 5c-d).

**Figure 5.**
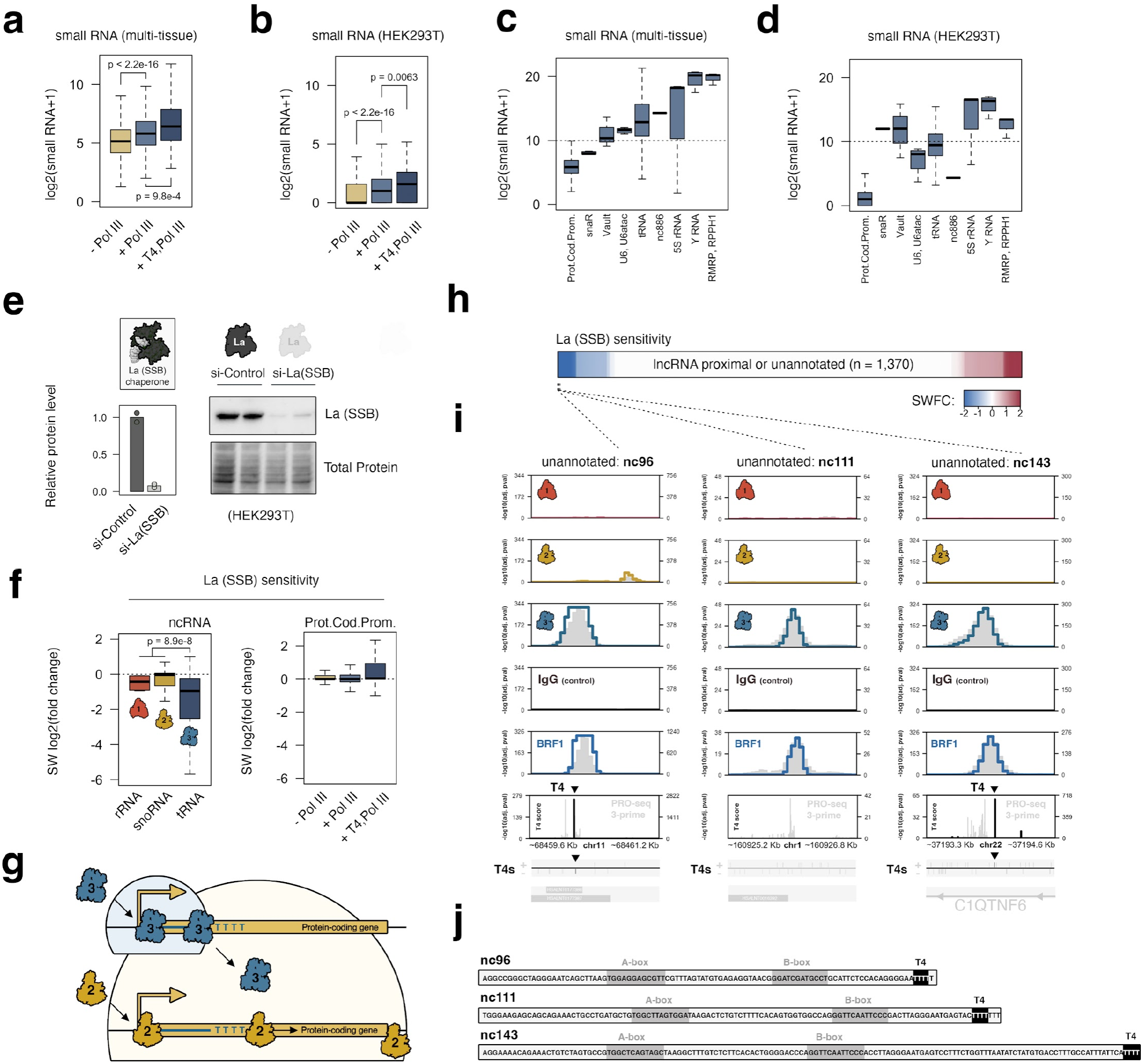
Depletion of RNA chaperone protein La (SSB) rules out the production of stable, promoter-derived small RNAs, but otherwise reveals multiple La-sensitive ncRNA species produced by RNA polymerase III at currently unannotated loci. **(a)** Distributions of multi-tissue small RNA levels mapped to Pol III-unoccupied (-Pol III), Pol III-occupied (+ Pol III), or Pol III-occupied protein-coding gene promoters with significant T4 enrichment (+T4, Pol III). **(b)** Analogous distributions of small RNA abundance linked to specific protein-coding promoters in HEK293T cells, which are sensitive to si-POLR3A and ML-60218 treatments. **(c)** Distributions of multi-tissue small RNA levels mapped to Pol III occupied protein-coding genes, compared to canonical Pol III-transcribed gene classes. **(d)** Analogous distributions of small RNA levels in HEK293T cells. **(e)** Immunoblot and quantification of RNA chaperone protein La following siRNA depletion in HEK293T cells. **(f)** Significance-weighted (SW) log2(fold change) of 45S pre-rRNAs, snoRNAs, and tRNAs following depletion of La (left), and of small RNAs mapped to protein-coding gene promoters of varying Pol III occupancy signatures. **(g)** Model: recruitment of Pol III to protein-coding genes results in transcription that terminates at T4 sequences, typically a short distance from the transcription start site, in contrast to processive transcription by Pol II. Pol III activity functionally enhances Pol II, but does not generate stable, promoter-derived small RNAs. **(h)** Heatmap visualization of La sensitivity for small RNAs from currently unannotated or lncRNA-proximal loci with significant Pol III signatures. **(i)** Examples of La-sensitive intervals with genomic indicators of Pol III activity. **(j)** Candidate sequence elements, related to 5i, on the basis of La-sensitivity, Pol III localization, and small RNA reads mapping to individual regions. Putative A-box, B-box, and termination sequence elements are highlighted.

Nevertheless, we further considered whether small RNA produced at specific protein-coding genes might depend on La (SSB), an RNA chaperone protein that plays critical roles in the stability, maturation, and transport of nascent Pol III-derived ncRNAs^36^. As an RNA-binding protein, La specifically recognizes the 3’-terminal oligo(U) tract that is universally present on these nascent ncRNAs, a feature driven by the termination mechanism of Pol III^13^. We reasoned that the presence of T4 termination and resulting 3’-oligo(U) tracts on nascent, promoter-derived small RNAs might similarly depend on La for RNA stability. To this end, we find that differential small RNA-seq analysis following La depletion confirms sensitivity of tRNAs to La disruption, in contrast to Pol I-derived 45S pre-rRNA and Pol II-derived snoRNAs (Figure 5e-f). However, examination of differential small RNA levels mapped to Pol III-occupied promoter windows does not indicate a similar pattern of dependency on La (Figure 5f). The minimal abundance and lack of Lasensitivity suggest that Pol III activity at protein-coding genes does not give rise to additional functional ncRNA species originating from these intervals. Instead, these results support a model in which Pol III recruitment and its transcription of protein-coding promoter sequences, constrained by T4 termination signals, is generally and primarily limited to enhancing the production of specific mRNA subpopulations (Figure 5g).

### Disruption of RNA chaperone protein La (SSB) uncovers potentially novel La-dependent Pol III-derived ncRNAs in humans

In addition to linking Pol III activity to protein-coding and other noncanonical genes, we finally considered whether our genomic atlas could successfully identify intervals in which Pol III transcription occurs beyond well-established coding, noncoding, and repeat sequences. We specifically queried genomic bins with significant Pol III signal and evidence of La-sensitivity, such that small RNA reads mapping to a given locus are significantly downregulated following La depletion. Out of 1,370 sites with Pol III occupancy that do not intersect any gene feature, or which otherwise intersect relatively uncharacterized lncRNAs^37^, we identify just a small number of regions with evidence of La-sensitivity (Figure 5h). Among the most compelling examples of putatively novel Pol III-derived ncRNAs, we show that strong Pol III occupancy and La-sensitivity converge on genomic windows proximal to uncharacterized lncRNAs on chromosomes 1 and 11, and to an unannotated locus within the protein-coding gene *C1QTNF6* on chromosome 22 (Figure 5i). Evidence of Pol III activity at these loci is further supported by significant BRF1 signals present across all three intervals, consistent with the canonical mechanisms of Pol III recruitment. In addition, significant T4 termination signatures are present at two of these sites, strongly supporting Pol III-driven production of nascent RNA (Figure 5i).

We infer the putative sequences of these ncRNAs (preliminarily named nc96, nc111, and nc142 on the basis of sequence length), by relying on the multi-context small RNA profiles generated across numerous human tissues and T4 termination signals (Figure 5j). Upon close examination of each putative ncRNA, we identify intervals with strong sequence similarity to both A-box and B-box gene-internal regulatory elements critical for TFIIIC recruitment to genes with type 2 promoter architectures^38^. Indeed, TFIIIC sub-unit GTF3C5 - a component of the DNA-binding TFIIIC2 subcomplex - is mapped to all three windows, consistent with established mechanisms of Pol III recruitment (Supplemental Figure 5)^39^. The confluence of sequence, recruitment, and post-transcriptional regulatory signatures, together, strongly support the classification of nc96, nc111, and nc142 as novel Pol III-derived ncRNAs for future investigation.

## Discussion

To date, limited efforts have been undertaken to understand the full extent of Pol III localization or the implications of Pol III activity beyond the boundaries of the currently established Pol III transcriptome. Our broad exploration of RNA polymerase occupancies, made possible by the 100s of ChIP-seq and RNA-seq datasets generated across human tissues and cell lines^35^, provides strong evidence of Pol III recruitment and transcription at specific protein-coding genes. These findings are supported by both Pol III occupancy and nascent RNA-seq analyses tailored to the unique termination mechanisms and resulting signatures tied to Pol III, and suggest that Pol III activity is limited to short elements constrained by the presence of T4 sequences capable of DNA-directed Pol III termination (Figure 5h). We note that previous Pol III transcription studies reported evidence of competent Pol III initiation at Pol II-transcribed gene promoters *in vitro*^40^, consistent with our discovery of widespread Pol III recruitment and transcription at Pol II-dependent protein-coding gene promoters in human cells. Though our work makes progress on understanding the significance of these events, our findings naturally give rise to new questions related to the mechanisms and implications of cross-talk between Pols II and III.

In the present study, a context-agnostic approach was made necessary by the currently limited breadth of Pol III mapping studies, precluding our ability to understand the contextual underpinnings of Pol III transcription at protein-coding genes. Nevertheless, if Pol III activity at protein-coding genes is restricted to specific cellular contexts or conditions, it may have been difficult to uncover Pol III activity at these sites without taking a comprehensive, multi-context perspective.

While current resources hint that BRF1, a TFIIIB complex subunit involved in Pol III recruitment at canonical type 1 and 2 promoters, is particularly enriched at Pol III-occupied protein-coding gene promoters, further delineation of the gene-specific Pol III recruitment mechanisms remain open-ended questions that future surveys of Pol III factors may address. Similarly, to what degree Pol III recruitment and activity at noncanonical loci is dependent on increased cellular levels and availability of Pol III, potentially driven under specific conditions, remains an important question for future research.

Functionally, the links between Pol III depletion and small molecule inhibition and the observed disruption (rather than enhancement) of mRNA produced from Pol III-occupied genes indicates that Pol III recruitment functionally promotes (rather than competes for) Pol II activity at these loci. The possibility of Pol III-dependent initiation and/or maintenance of an active chromatin environment, perhaps also intersecting promoter-proximal pausing mechanisms, are tempting preliminary models of speculation. In either case, the biological significance of this effect appears to be tied to protein-coding genes central to biosynthesis and cellular growth. We show, for example, that Pol III is recruited to numerous genes encoding ribosomal proteins and, additionally, that high expression of Pol III-occupied and Pol III-sensitive genes is linked to unfavorable outcomes in cancer. These findings point to a novel form of cross-talk between Pols II and III, such that increaseing Pol III activity may promote complementary increases in ribosomal proteins and other growth-related factors.

Finally, our study suggests that the small ncRNA products produced by Pol III at protein-coding gene promoters are minimally abundant, potentially due to rapid turnover. Although this result limits the notion of novel functional small RNA species derived from Pol III activity at these loci, we nevertheless uncover evidence for novel small RNA species derived from other genomic intervals that currently lack clear gene annotation features. While the significance of these ncRNAs will require targeted functional genomic studies, these data altogether reveal new and unexpected insights into the breadth of Pol III transcription.

## Limitations of this study

Due to the context-agnostic nature of our RNA polymerase metamap, we are unable to define the specific cellular contexts in which Pol III activity is tied to a particular gene. However, our complementary Pol III disruption experiments (si-POLR3A and ML-60218) demonstrate a link between Pol III and Pol III-occupied protein-coding genes specifically in HEK293T, a transformed human embryonic kidney cell line amenable to functional studies. Application of large-scale, integrative genomic approaches targeting Pol III in a variety of cellular contexts is therefore needed to fully appreciate and delineate the dynamic range of Pol III expansion under specific contexts and conditions.

We additionally note that our pursuit of establishing Pol III occupancy patterns is dependent on the most recent “snapshot” of gene annotations in humans, and that several non-trivial decisions, such as setting gene-class priorities, gene promoter window sizes, and defining which genomic intervals to mask, play an important role in shaping the outcome of our study. In this way, numerous examples of Pol III occupancy may be missed in our metamap - a trade-off between sensitivity and specificity. Thus, potentially promising but uncalled Pol III target genes (i.e. false negatives) is an additional limitation of our study.

## Methods

### Cell culture and RNAi / small molecule Pol III inhibition

HEK293Tcells were obtained from ATCC (CRL-3216, Batch#70049877) and grown in 10 cm BioLite™ Cell Culture Treated Dishes (Catalog# 12-556-002 Thermo Scientific) in Dulbecco’s Modified Eagle Medium, high glucose (Catalog# 11965092, Gibco). HEK293T experiments were conducted on cells between passage 10-25.

For POLR3A depletion experiments, HEK293T cells were seeded into 6 well Nunc™ Cell-CultureTreated Multidishes (Thermo Scientific, Catalog# 140675) and incubated overnight. siRNA (listed below, IDT) is transfected using Lipofectamine® RNAiMax (cat # 13778150). Cells were incubated for 48 hours at 37 °C with 5% CO2 before collection. 80% of cell pellets are used for RNA extraction, performed using E.Z.N.A.® Total RNA Kit I (Catalog# R6834-01, Omega), with 100% ethanol instead of 70% at the binding step to enrich small RNA. 20% cells are saved for western blot.

|  |  |  |
| --- | --- | --- |
| Scramble siRNA | Silencer™ Select Negative Control No. 1 siRNA (Catalog# 439084, Invitrogen™) | 20nM |
| POLR3A siRNA | rCrCrUrGrCrArGrCrUrArUrGrGrArGrGrGrArATT<br>rUrUrCrCrCrUrCrCrArUrArGrCrUrGrCrArGrGTT | 20nM |

For La (SSB) depletion experiments, HEK293T cells were seeded into 12-well Cell-CultureTreated Multidishes (Thermo Scientific) and incubated overnight. HEK293T cells were then transfected with 150nM siRNA using Lipofectamine® RNAi Max (cat # 13778150) for 48 hours at 37 °C with 5% CO2 before collection. siLa/SSB (IDT): rArGrA rUrUrG rGrArU rGrCrU rUrGrC rUrGrA rATT; si-scramble (Invitrogen): Silencer Select Negative Control #1 siRNA. RNA extraction was performed using mirVana PARIS RNA and Native Protein Purification kit according to the manufacturer’s instructions (Invitrogen).

For Pol III inhibition, ML-60218 (MedChemExpress, HY-122122) was added to 80% confluent HEK293T cells to a final concentration of 25uM, and cells are collected 4h post treatment.

### Reverse Transcription qPCR

For small RNA qPCR following POLR3A depletion, cDNA was first generated using the TaqMan MicroRNA Reverse Transcription Kit (Catalog # 4366597, ThermoFisher). qPCR was then performed using TaqMan™ Fast Advanced Master Mix (Catalog#4444557, Applied Biosystems) and TaqMan™ MicroRNA Assay (Catalog # 4440887, 4398987, thermo fisher)

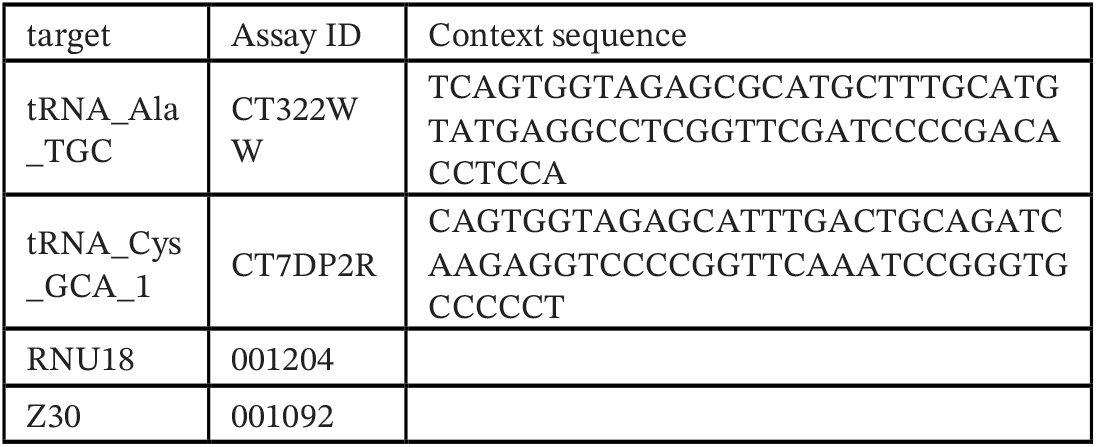

### Immunoblotting and protein quantification

Cell pellets were washed once with PBS before lysis with RIPA buffer (Catalog#J62524.AD, Thermo Scientific) following standard protocols. Total protein concentration was determined using Pierce BCA protein assay kit (Catalog# 23225, Thermo Scientific), and equivalent protein fractions were diluted in cell lysis buffe, and 4*Laemmli Sample Buffer (Catalog#1610741, BIO-RAD). Proteins were separated on 4–20% Mini-PROTEAN® TGX Stain-Free™ Protein Gels, 15 wells (Catalog#4568096, BIO-RAD) using 10x Tris/Glycine/SDS (Catalog#1610732, BIO-RAD) and transferred onto polyvinylidene difluoride membranes 0.2 um (Catalog# LC2002, Invitrogen) with Trans-Blot® Turbo™ Transfer System (Catalog# 1704150, BIO-RAD). Transfer membranes were blocked with 5% blotting-grade blocker (Catalog# 1706404, BIO-RAD), followed by incubation with the primary antibodies at 4°C overnight. Membranes were washed with TBST and incubated with mouse and rabbit secondary antibodies conjugated with horseradish peroxidase (Catalog# 31430, 31462, Invitrogen) for 2 h at room temperature followed by three washes in TBST. Proteins were visualized using Super SignalWest Pico (Catalog# 34580, Thermo Scientific) or SuperSignal WestFemto (Catalog#34096, ThermoScientific) with ChemiDoc™ Touch Imaging System (Catalog# 1708370, BIO-RAD). Protein abundances were calculated by normalizing to reference protein Lamin B2 or β-tubulin and to total protein.

Antibodies:

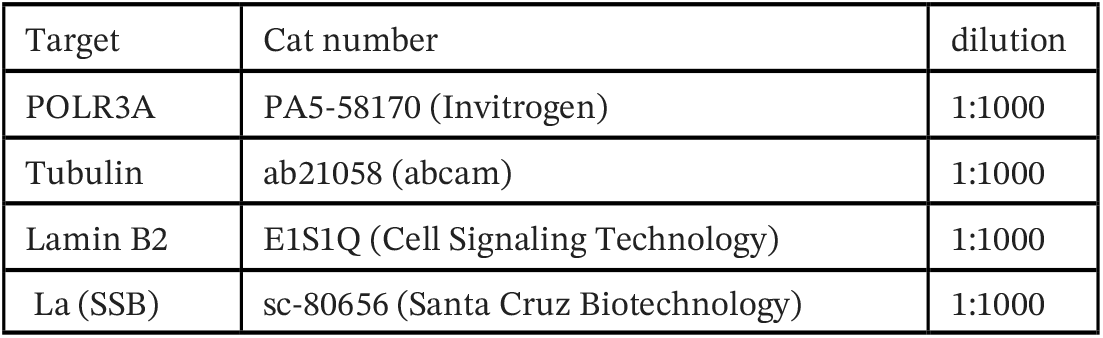

### DNA library preparation and sequencing

For RNA-seq experiments following POLR3A depletion and Pol III inhibition, libraries were prepared with the Kapa Hyper Stranded mRNA library kit (Roche) and sequenced on two 10B lanes for 151 cycles from both ends of the fragments on a NovaSeq X Plus with V1.0 sequencing kits. For small RNA-seq experiments following La (SSB) depletion, samples were first treated with Antarctic Phosphatase (NEB), followed by treatment with T4 Polynucleotide Kinase (NEB) according to the manufacturer’s protocol. Small RNA libraries were generated using the NEBnext small RNA library preparation kit according to instructions (New England BioLabs). small RNA high-throughput sequencing libraries were sequenced for 101 cycles using NovaSeq X Plus. All sequencing was performed at the University of Illinois, Roy J.Carver Biotechnology Center. Fastq files were generated and demultiplexed with the bcl2fastq v2.20 Conversion Software (Illumina).

### RNA-seq alignment and differential analysis

RNA-seq reads were first trimmed using TrimGalore^43^ and then mapped to the hg38 transcriptome and quantified with Salmon^51^. Differentially expressed genes were determined using DESeq2^52^. small RNA-seq reads were first trimmed using TrimGalore^43^ and then mapped to the entire genome space (GRCh38) using bowtie2^44^. By default, multi-mapping reads were reported as a singular best alignment. Promoter-specific raw small RNA signal counts were extracted using bedtools^48^. Differentially expressed small RNA was determined using edgeR^58^.

### Multi-class promoter curation

Protein-coding transcript annotations were retrieved from Ensembl (Gencode V43)^41^, whereas annotations for noncoding gene types were retrieved from RNAcentral (V22)^42^. For each URSID in RNAcentral, we concatenated the gene information across over 30 databases linked to RNAcentral. Clustering of TSS was performed to eliminate redundancy of TSS within each gene type, with clusters identified within a 120-nucleotide vicinity for each gene type and strand. Gene information pertaining to TSSs of the same cluster are collapsed together. Promoter regions were defined based on the position of the TSS cluster median, spanning from 50 nucleotides upstream to 150 nucleotides downstream. Repeat annotations were retrieved from RepeatMasker. Finally, all promoter windows associated with “blacklisted” genomic intervals (ENCFF356LFX) were masked from further analysis. The workflow and resources related to the Multi-class promoter curation method are available at https://github.com/VanBortleLab/multiClass PromoterCuration.

### Uniform RNA polymerase occupancy scoring

We leveraged ChIP-seq data from GEO^46^ and ENCODE^35^, encompassing 2, 153, and 47 datasets for Pol I, Pol II, and Pol III, respectively (Supplemental Table 1), with corresponding total mapped reads of 53 million, 7 billion, and 5.6 billion. We implemented a comprehensive normalization procedure: first, in the context of Pol III and Pol II, reads were normalized across each context for every available subunit (including POLR3A, POLR3B, POLR3C, POLR3D, POLR3E, POLR3G and POLR2A), ensuring parity in signal representation across each cellular context. Subsequently, in the context of Pol III, we normalized reads for each subunit to equalize their contributions, thus preventing biases stemming from variable sequencing depths of subunits. After normalizing subunits of Pol III, we computed the median occupancy signal across all Pol III subunits. In the context of Pol I, we chose to concatenate reads from POLR1A and POLR1B, the only two experiments currently available in humans. Finally, we scaled each polymerase to 100 million reads ensuring consistency in signal intensities and enabling meaningful comparisons between polymerases. This process facilitated unbiased analysis of RNA polymerase occupancy and dominance across the genome.

Following read normalization, the signal coverage for each polymerase was extracted across binned intervals of 50 bp. Enrichment scores for each bin were determined using a Poisson framework. Briefly, for each bin, the expected signal within 5 Kb (λ5k), 10 Kb (λ10k), and across the entire genome (λgenome) is computed. The λ value is thereafter determined as the maximum lambda value of the distance set. The probability density function (PDF) utilized is given by P(X = x) = (e^(-λ) * λ^x) / x!. If k represents the observed signal for the bin, the p-value tied to a specific bin enrichment is calculated using P(X > k). P-values were adjusted globally using the Benjamini-Hochberg method. Finally, an enrichment score is computed as the negative logarithm (base 10) of the adjusted p-value. In cases where infinite scores are encountered, they are assigned the highest non-infinite score. The workflow and processed data are available at https://github.com/VanBortleLab/RNApolymerase_metamap.

### RNA polymerase dominance visualization

Initially, RNA polymerase scores were assigned to each gene promoter by selecting the maximum score from the bins intersecting the promoter window. Subsequently, RNA polymerase dominance plots were generated by representing the three RNA polymerase scores for each gene as three point masses set at three vertices of an equilateral triangle, and plotting the center of mass of this system. getCOM() and plotTriangle() functions, which calculate and plot the center of mass for each gene, respectively, are available at https://github.com/VanBortleLab/dominatR.

### Nascent T4 score calculation

We retrieved all available nascent PRO-seq (Precision Run-on Sequencing^22^) datasets available via ENCODE^35^ corresponding to 18 distinct experiments across 7 different cell lines. To ascertain strand specificity, we employed the infer_experiment code from the RSeQC^47^ package and filtered out 5 experiments with a strand specificity of less than 85%, resulting in 13 experiments (Supplemental Table 2). Subsequently, we utilized genomecov from BEDtools^48^ to create the 3’ pileup of reads across the genome, employing the flags “-dz -3”. T4 motifs (TTTT) were identified within the hg38 genome using SeqKit^49^, with a 3-nt cushion applied on either side of the T4 motif, resulting in a 10-nt long T4 motif. Employing the same scoring method utilized for the occupancy score of 50-nt bins, we computed the T4 score, this time focusing on the 10-nt T4 motif with expected signal over two vicinities: 20 nt and 50 nt on either side of the T4 motif. The workflow and processed data are available at https://github.com/VanBortleLab/NascentT4score.

To test the significance of enrichment of T4 scores in pol3-enriched active genes compared to pol2-enriched active genes, we conducted a significance test using random sampling. Specifically, we generated a distribution of T4 scores by summing the T4 scores of genes randomly sampled from the list of pol2-enriched genes. The number of genes thus sampled from the pol2-enriched set equaled the number of genes in the pol3-enriched set. This sampling process was repeated 1 million times to create a robust distribution of T4 scores. The p-value was determined as the fraction of these 1 million sums greater than the sum of T4 scores of our pol3-enriched genes. This approach allowed us to assess the significance of T4 score enrichment in pol3-enriched active genes relative to pol2-enriched active genes.

### ChIP-seq Transcription Factor (TF) enrichment analysis

Protein coding genes were divided into two groups: one comprising 1572 genes with a median pol3 score greater than or equal to 2, and the other consisting of genes failing to meet this threshold. Subsequently, utilizing the ChIPatlas enrichment tool ^24^, a list of transcription factors (TFs) across multiple experiments and cell types, along with their enrichment (-logqval and FoldEnrichment (FE) value) on the first group of genes compared to the second group was found. Experiments with fewer than 5000 peaks were filtered out. Enrichment scores were calculated using the formula SCORE = (-logqval) * log2(FE). For each TF, scores were collapsed within the same cell type using the mean score, and then across different cell types using the median score.

### Multi-tissue small RNA-seq abundance

We retrieved a total of 204 small RNA-seq datasets (Supplemental table 3) available via ENCODE^35^ derived from 64 distinct contexts (i.e. cell lines and tissue types). Small RNA read pile-ups were determined using multiBamSummary from deeptools^50^ at 10 bp resolution. Reads were aggregated by cellular context and a summary RNA profile was thereafter determined by scaling all context-specific RNA reads to the minimal sequencing depth observed across all 64 contexts. Multi-tissue promoter-linked small RNA levels were then determined as the sum of normalized signals across bins within 100 bp of each promoter window.

### TCGA gene correlation and survival features

The Cancer Genome Atlas (TCGA) primary solid tumor gene expression levels were extracted across all disease cohorts from the Broad Institute TCGA Genome Data Analysis Center (GDAC) Firehose mRNASeq Level 3 RSEM gene normalized data files at https://gdac.broadinstitute.org. Individual pan-cancer Pearson correlation scores were calculated for each Pol III-specific subunit against all genes included in these data. The strength and direction of correlation signatures related to Pol III-occupied and Pol III-sensitive genes were then measured by comparing the observed distribution of correlation metrics of a given gene set against an empirical null distribution derived by 10,000 permutations. Survival statistics linking gene expression levels to patient outcomes were retrieved from tcga-survival.com^53,54^.

### Functional enrichment analysis

Gene Ontology enrichment analysis was performed using the R package clusterProfiler^55,56^ with genome wide annotation for Human org.Hs.eg.db^57^. Gene sets containing those associated with Pol III occupancy or Pol III sensitivity were compared against a universe of genes restricted to those with RNA polymerase occupancy (set 1) or inclusion in differential analysis (i.e. present following low count filtering, set 2). Enrichments were calculated for GO cellular component and GO biological processes and p values adjusted for multiple hypothesis testing.

### Identification of unannotated, putative Pol III products

To identify unannotated putative Pol III products, at first, genomic bins with a median Pol3 score less than 2 were removed. Next, bins within 350 base pairs of any known annotations’ transcription start sites (TSS) were removed. In cases of consecutive adjacent bins, only the bin with the maximum median Pol3 score was chosen. Finally, bins within blacklisted regions were excluded, and the resulting bins were extended by 75 base pairs on each side to generate a list of 659 unannotated putative Pol3 promoters.

## Resource Availability

The mRNA-seq and small RNA-seq data generated for this study are available through the NCBI Gene Expression Omnibus with accession numbers GSE268457 and GSE268497, respectively.

## Acknowledgements

We thank Alvaro Hernandez, Chris Wright, Danman Zhang, and staff at the Carver Biotechnology Center for sequencing services, and administrators of the Carl R. Woese Institute for Genomic Biology (UIUC) Biocluster for computational support. We thank Prof. Brian Freeman, Prof. Andy Belmont, and members of the Van Bortle lab for critical reading of the manuscript and helpful suggestions. This work was supported by the National Institutes of Health, National Human Genome Research Institute (NHGRI) grant R00HG010362 to K.V.B.

## Author contributions

Study design: RKC, RC, SZ, SL, DS, KVB. Data collection: RKC, RC, SZ, KVB. Data analysis: RKC, RC, SZ, KVB. Data interpretation: RKC, RC, SZ, KVB. Writing: RKC, RC, SZ and KVB with comments from all authors.

## Supplemental Figures

**Supplementary Figure 1.**
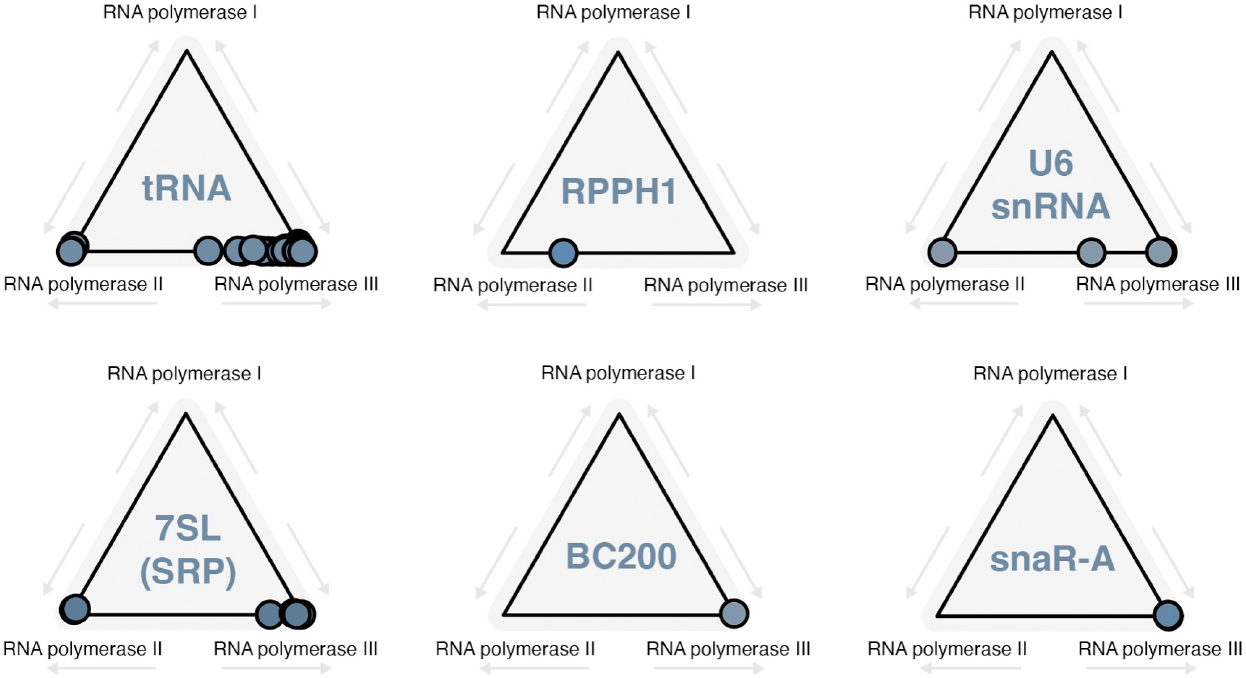
RNA polymerase dominance at subtypes of canonical Pol III-transcribed genes (related to Figure 2A). RNA polymerase dominance plots for canonical Pol III-transcribed small noncoding RNA genes, including genes encoding tRNAs, *RPPH1*, U6 snRNA, 7SL signal recognition particle (SRP) RNA, Brain Cytoplasmic RNA 200 (BC200, *BCYRN1*), and small NF90-associated RNA isoform A (snaR-A).

**Supplementary Figure 2.**
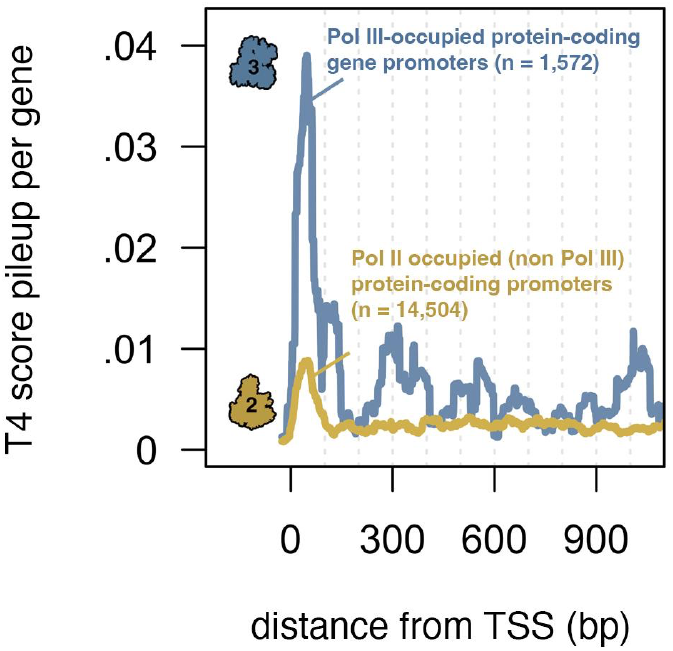
T4 score pileup as a function of intragenic T4 distance from protein-coding gene promoters. Line plots represent the composite magnitude of T4 scores by intragenic distance from a protein-coding gene promoter start site. Scores corresponding to each T4 motif occurrence at a specified distance were taken in aggregate. Pileup signal plots were thereafter smoothed using a 50-nt sliding window. Comparison of protein-coding gene promoters bound by Pol III (blue) with protein-coding gene promoters unbound by Pol III (gold).

**Supplementary Figure 3.**
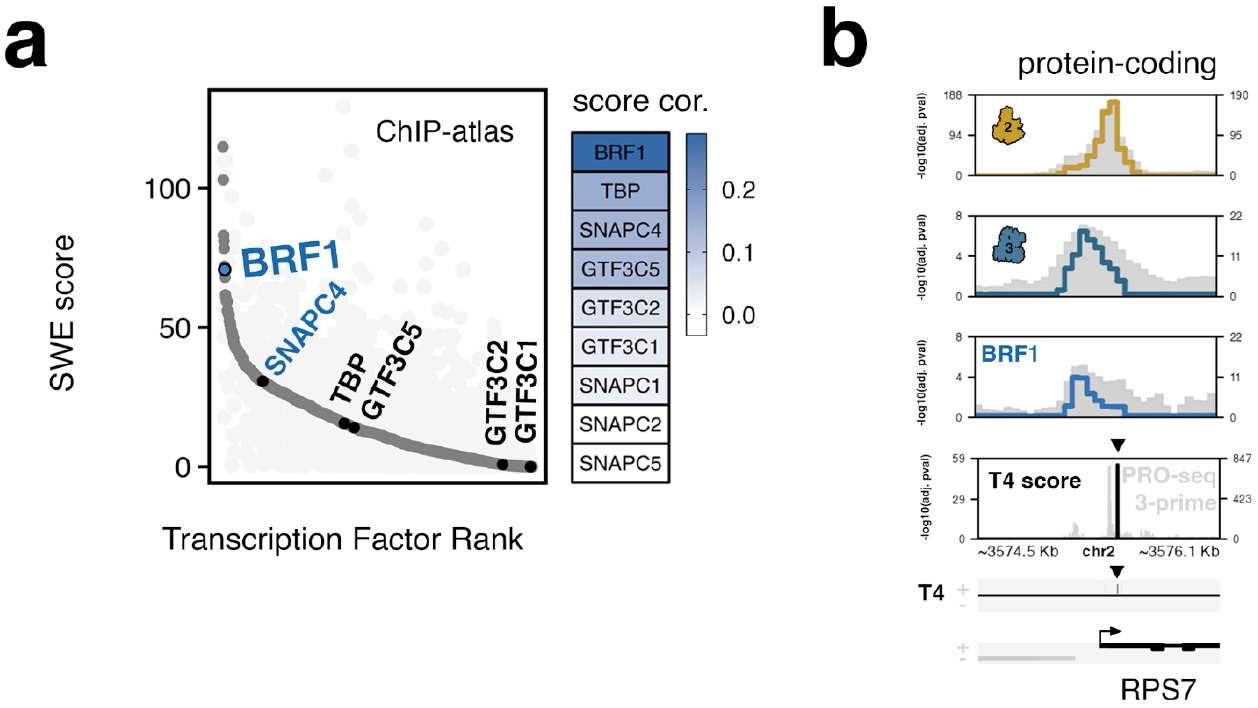
Transcription factor overlap enrichment analysis at Pol III-occupied protein-coding gene promoters. **(a)** Ranked overlap enrichments for individual transcription factors previously mapped by ChIP-seq. Established Pol III transcription factor-related proteins are highlighted. (BRF1 is a subunit of the TFIIIB transcription factor complex). **(b)** Visualization of significant Pol III and BRF1 occupancy scores at RPS7, an example protein-coding gene encoding Ribosomal Protein S7

**Supplementary Figure 4.**
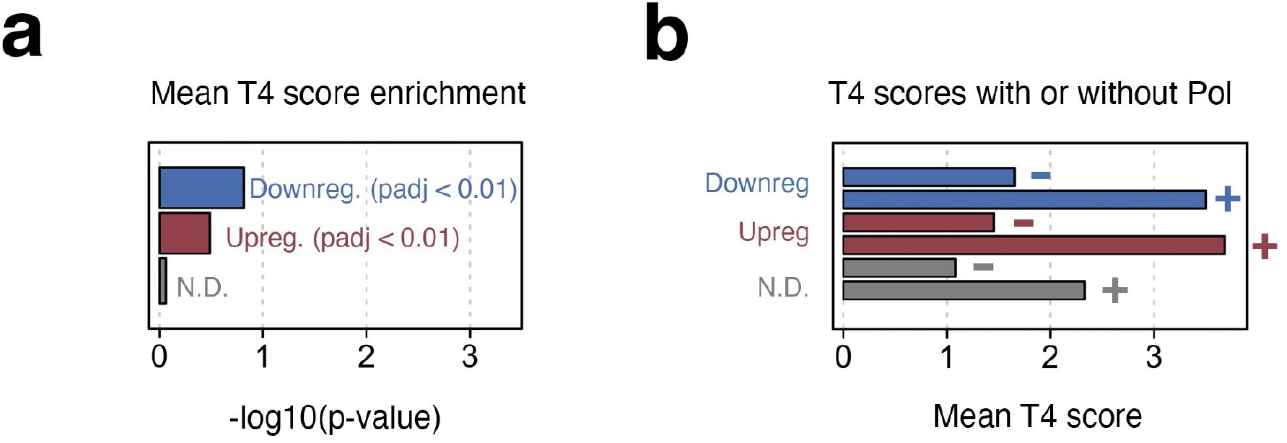
Enrichment of nascent T4 scores at shared si-POLR3A and ML-60218 sensitive genes. **(a)** Enrichment analysis of mean T4 scores at down-, up-, and nondifferential (N.D.) gene sets (related to Figure 3i). **(b)** Mean T4 scores for genes marked by significant Pol III signal (+) compared to genes lacking significant Pol III signal (-), at down-, up-, and nondifferential (N.D.) gene sets.

**Supplementary Figure 5.**
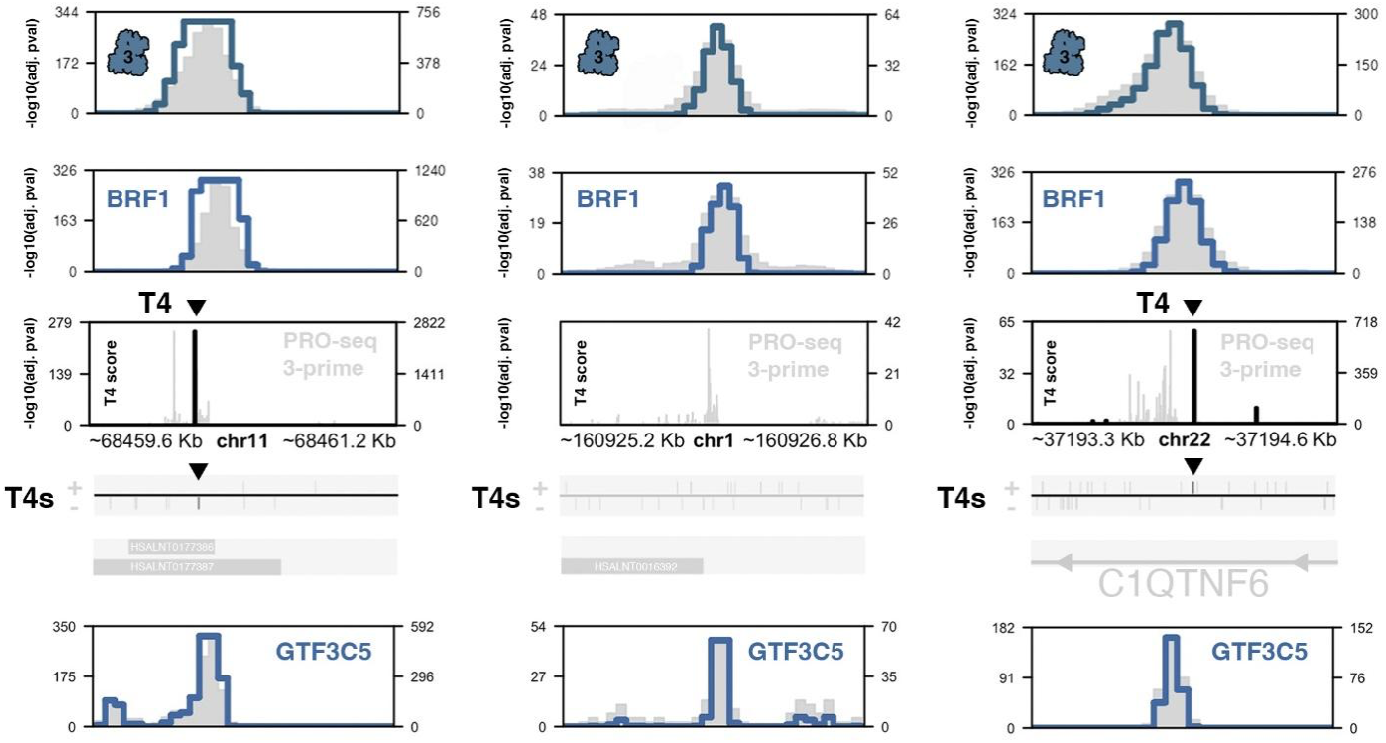
GTF3C5, a DNA-binding component of the TFIIIC complex, is present at putatively novel Pol III-transcribed gene loci. Signal plots for Pol III, BRF1, nascent T4 signatures at unannotated genomic intervals associated with nc96, nc111, and nc143 (related to Figure 5i), with the addition of TFIIIC subunit GTF3C5 (bottom).

## Supplemental Tables

**Supplementary Table 1.**
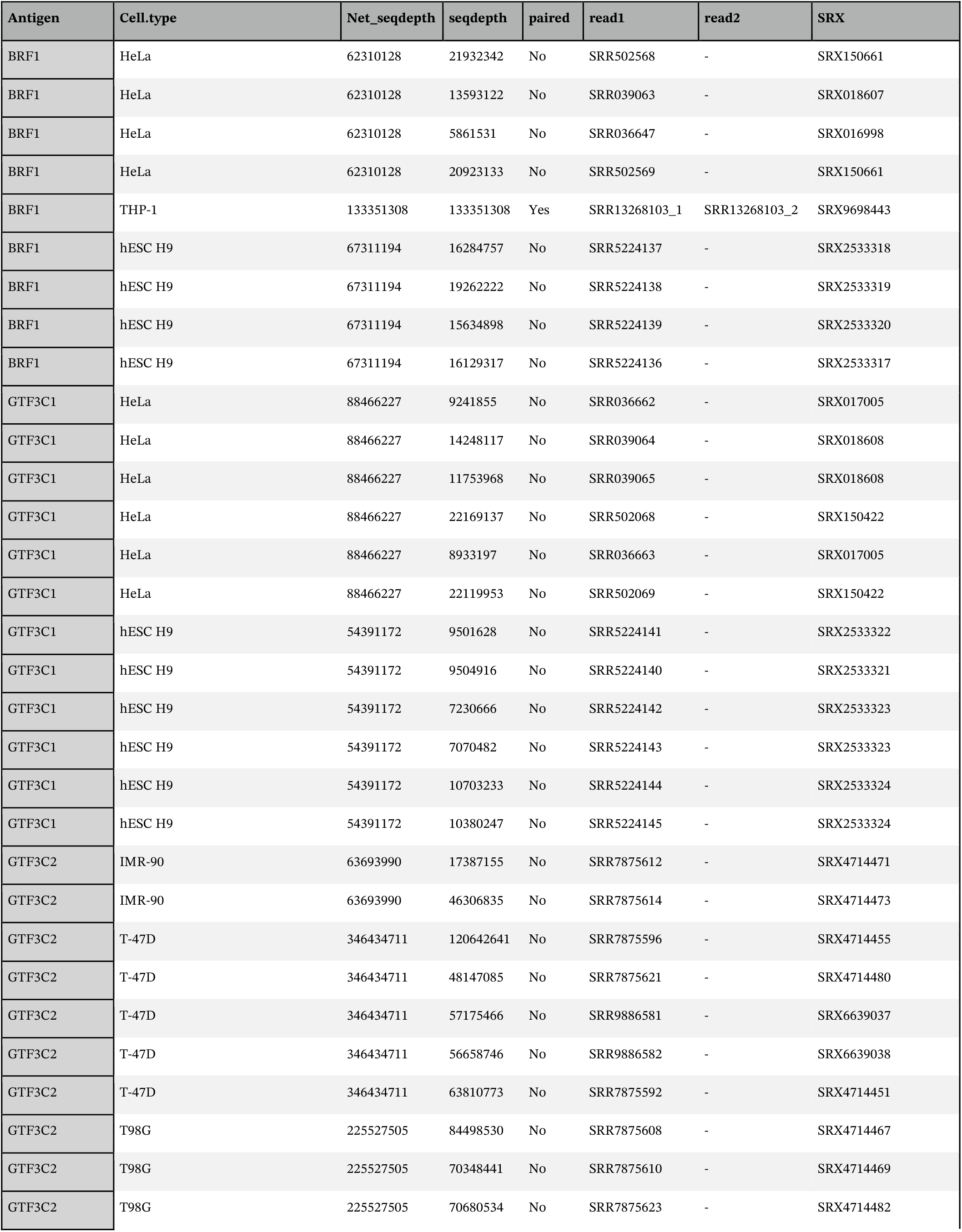
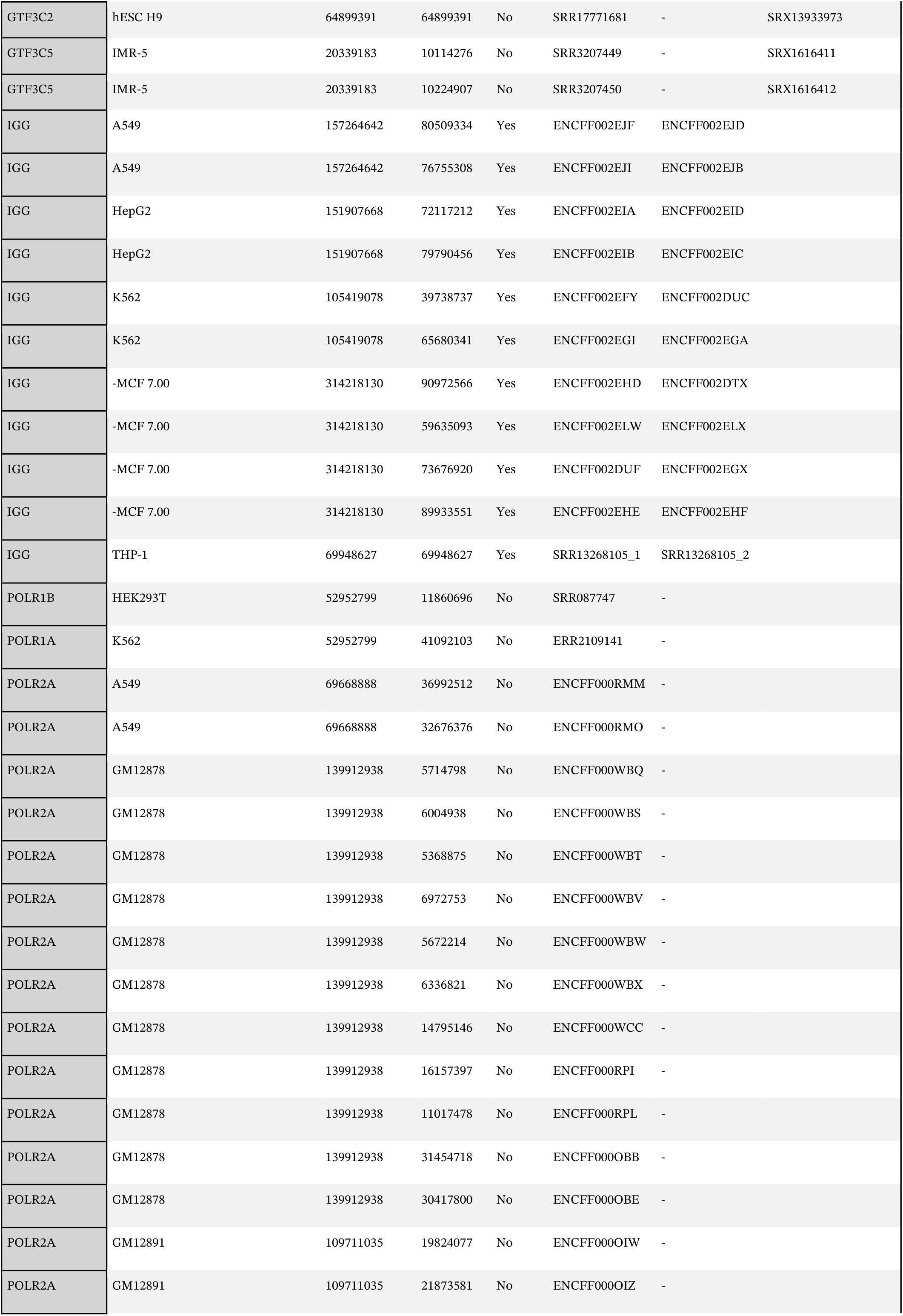
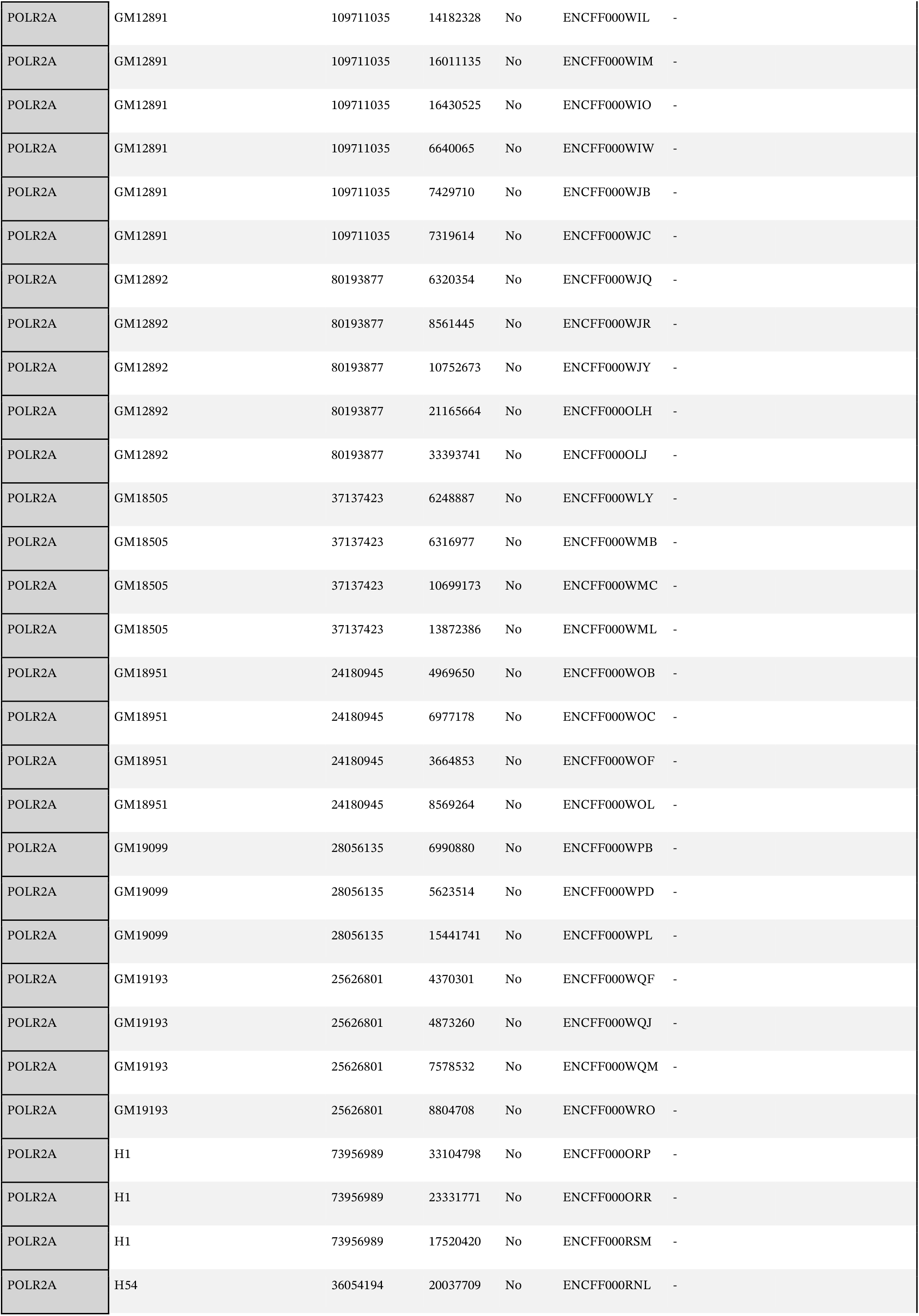
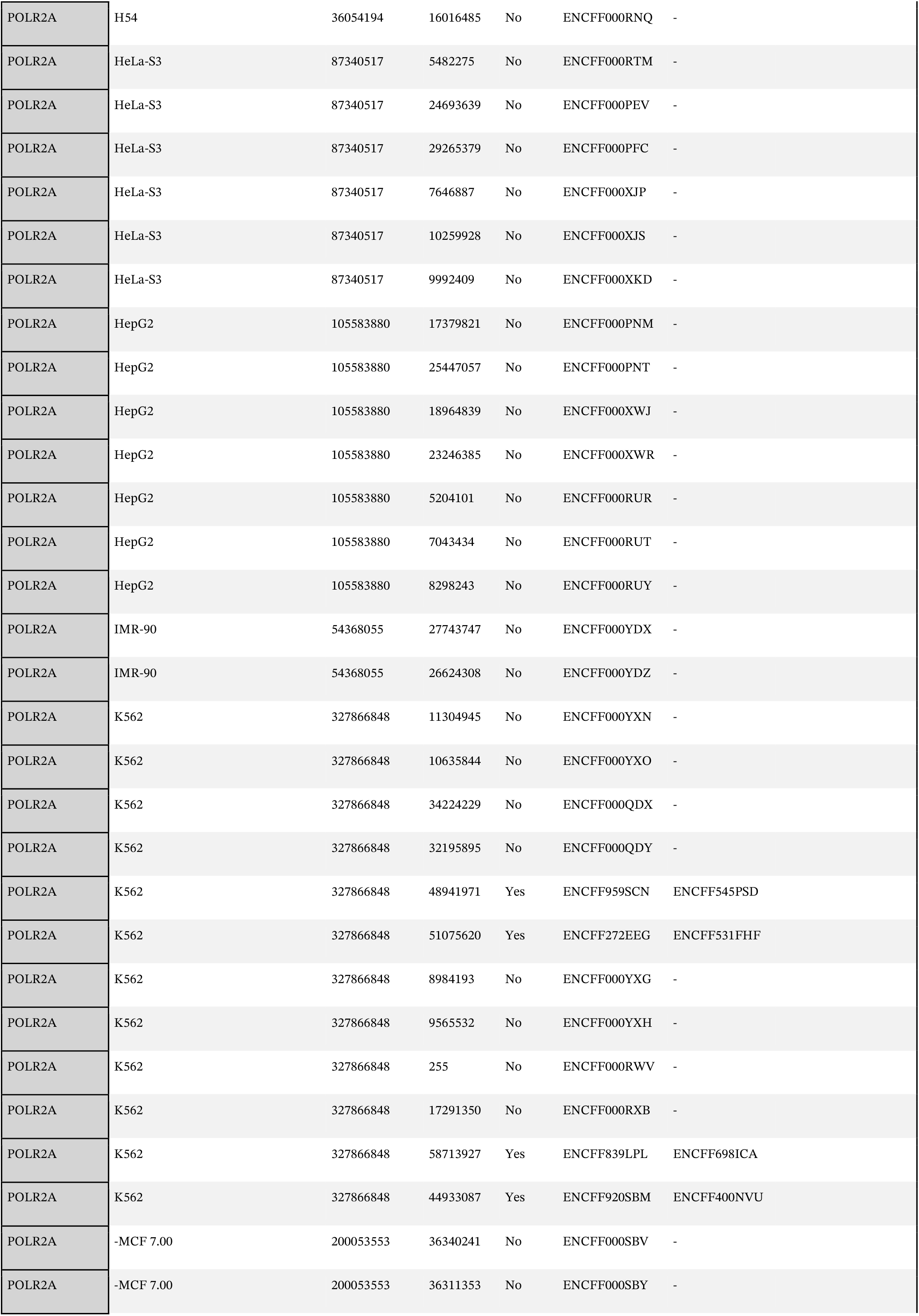
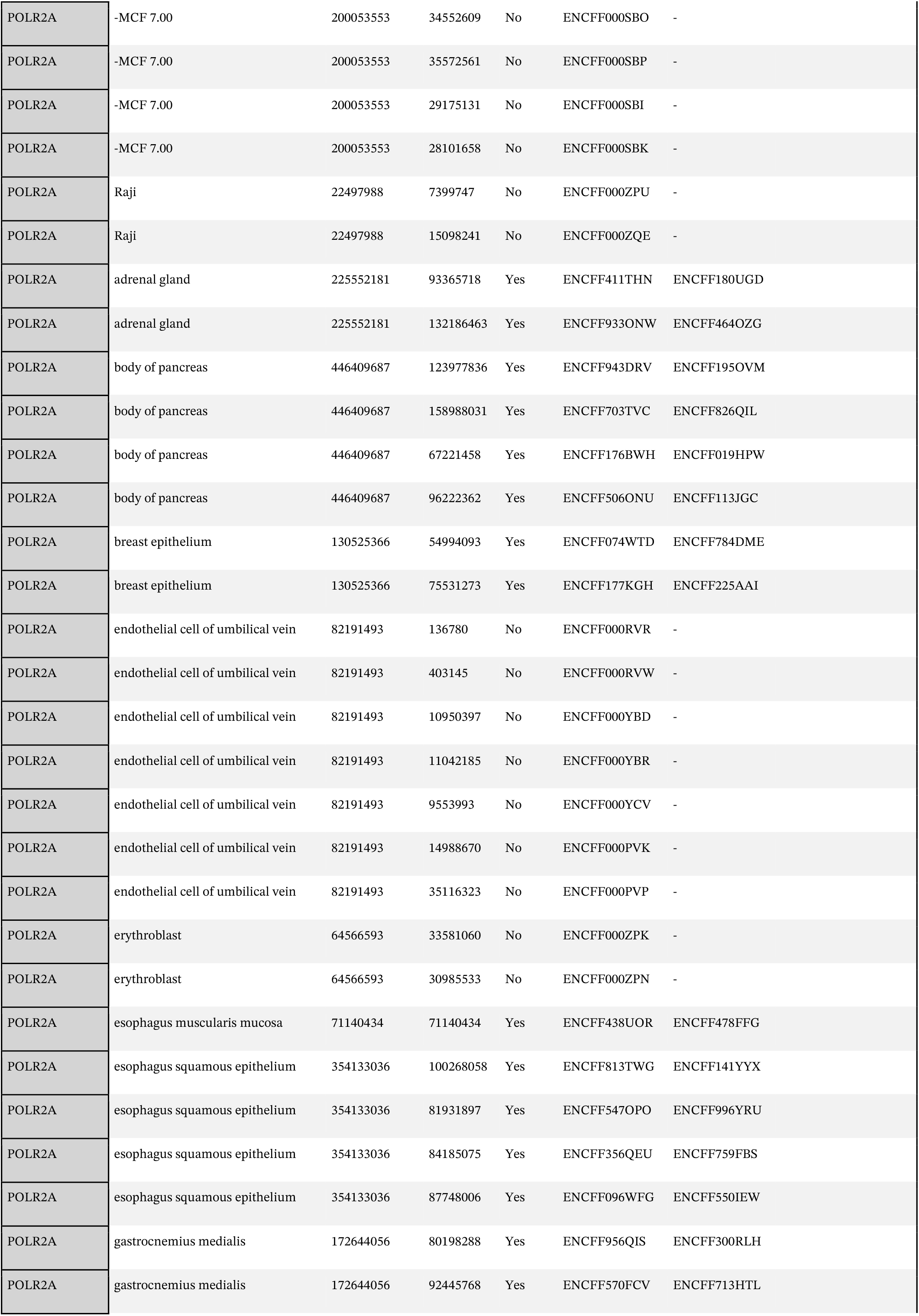
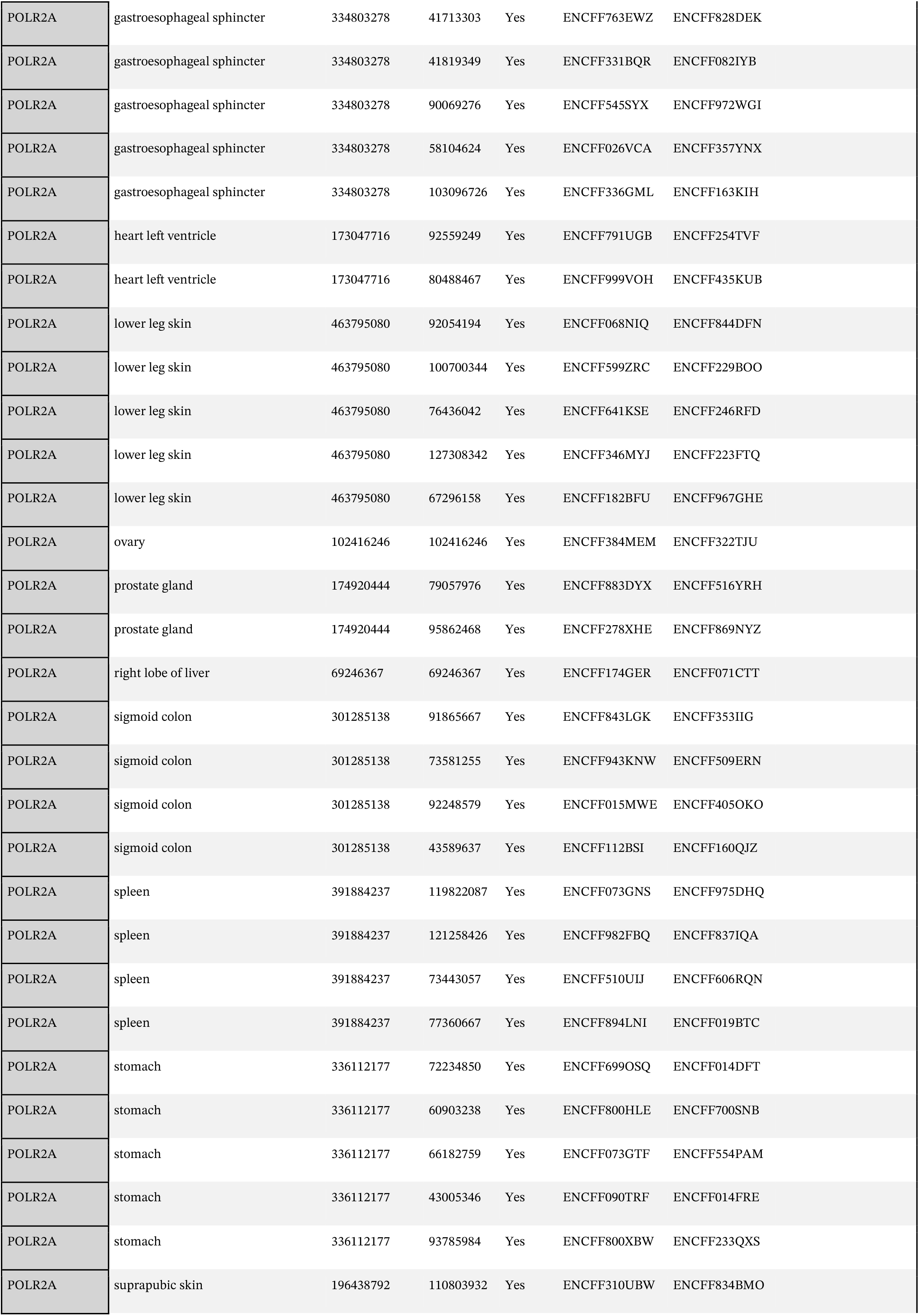
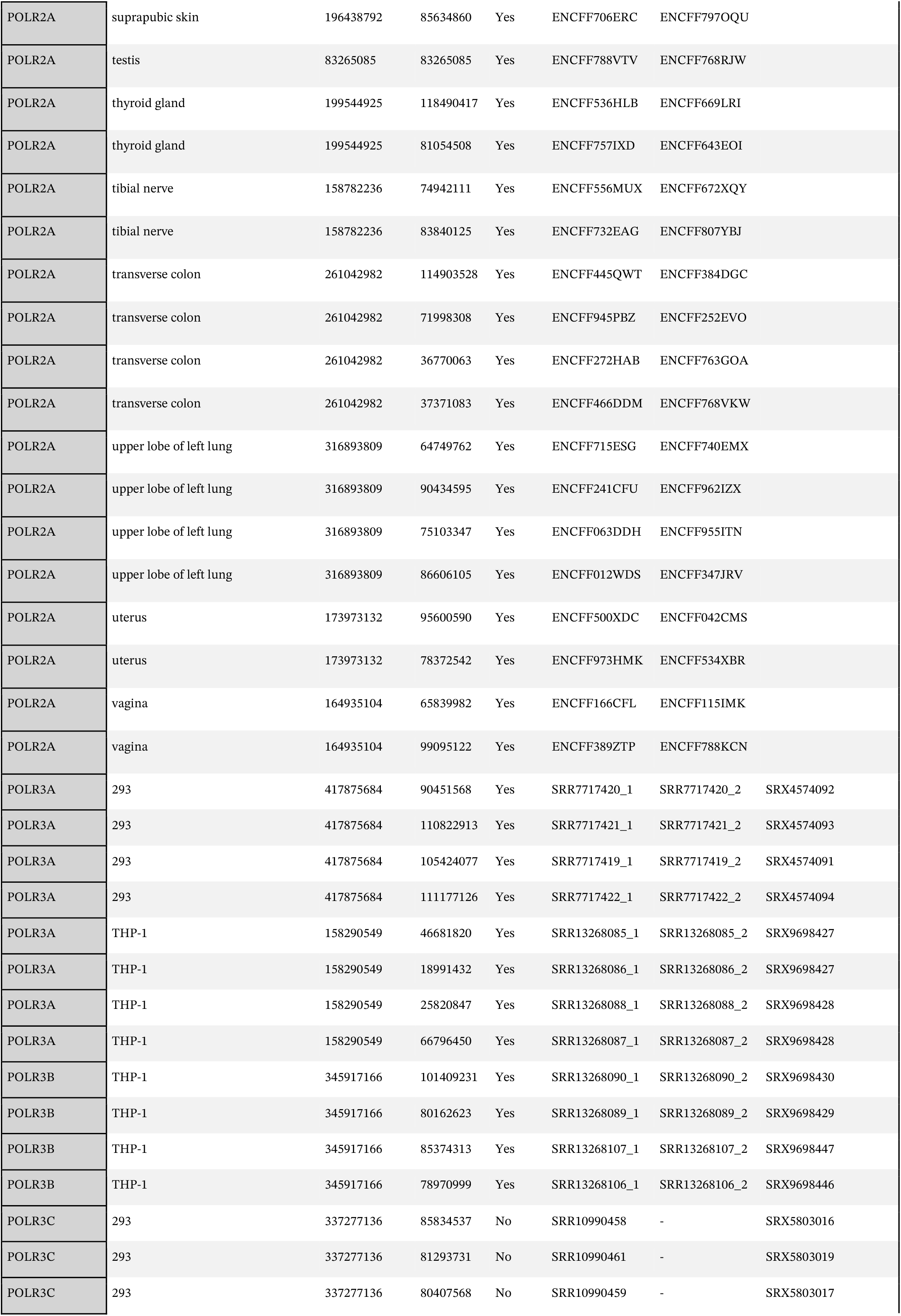
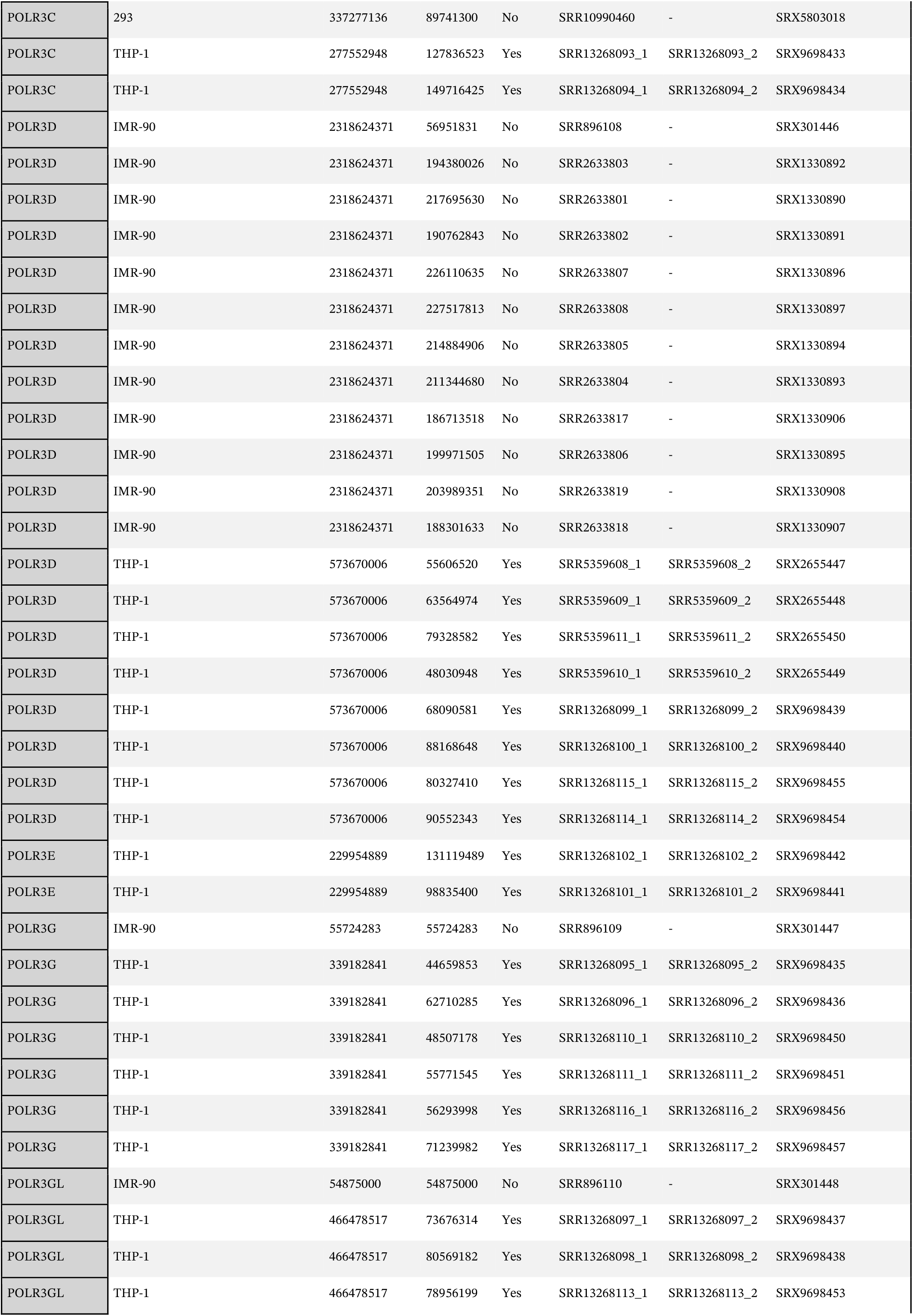
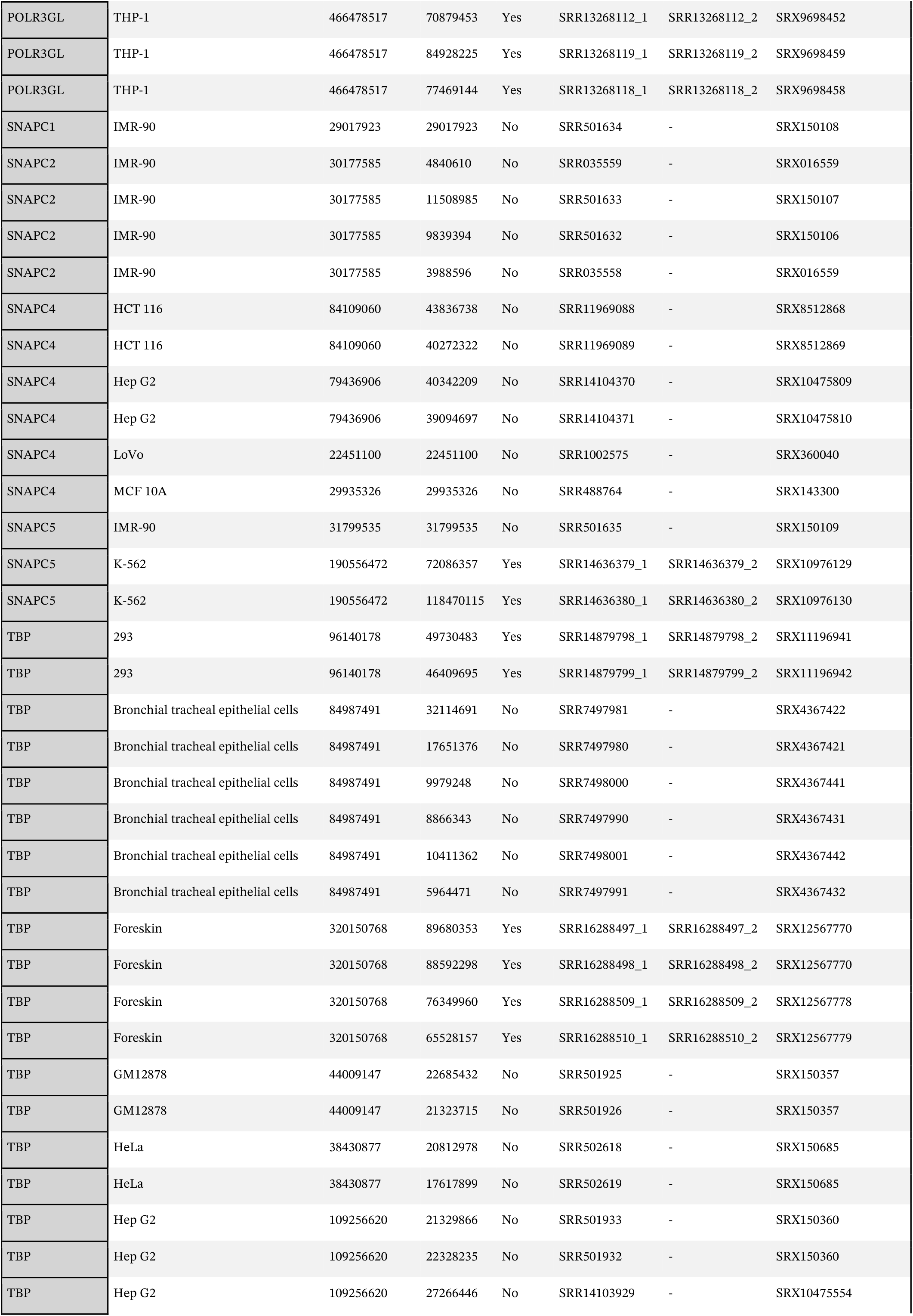
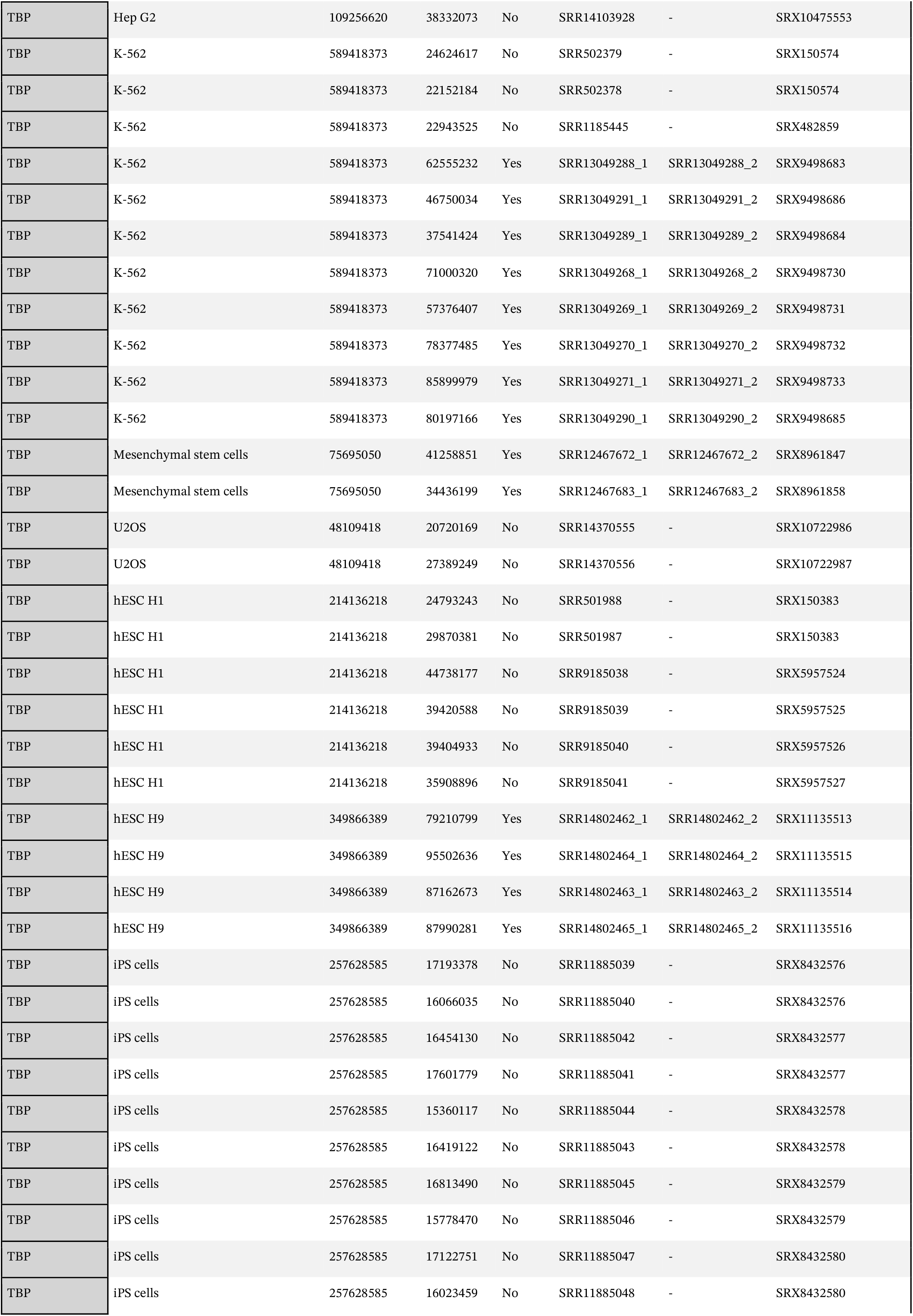
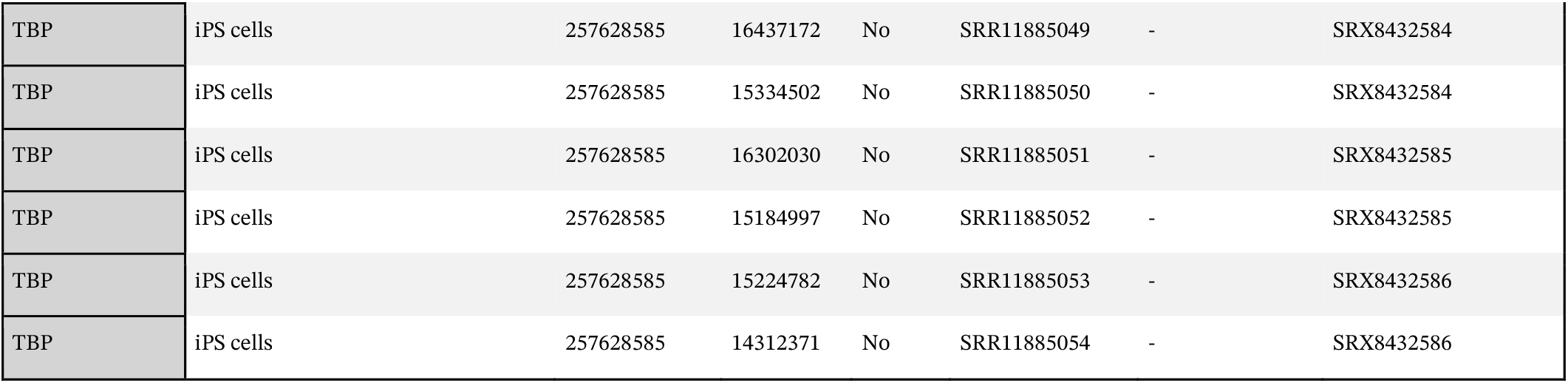
ChIP-seq data used in this study.

| Antigen | Cell.type | Net_seqdepth | seqdepth | paired | read1 | read2 | SRX |
| --- | --- | --- | --- | --- | --- | --- | --- |
| BRF1 | HeLa | 62310128 | 21932342 | No | SRR502568 | - | SRX150661 |
| BRF1 | HeLa | 62310128 | 13593122 | No | SRR039063 | - | SRX018607 |
| BRF1 | HeLa | 62310128 | 5861531 | No | SRR036647 | - | SRX016998 |
| BRF1 | HeLa | 62310128 | 20923133 | No | SRR502569 | - | SRX150661 |
| BRF1 | THP-1 | 133351308 | 133351308 | Yes | SRR13268103_1 | SRR13268103_2 | SRX9698443 |
| BRF1 | hESC H9 | 67311194 | 16284757 | No | SRR5224137 | - | SRX2533318 |
| BRF1 | hESC H9 | 67311194 | 19262222 | No | SRR5224138 | - | SRX2533319 |
| BRF1 | hESC H9 | 67311194 | 15634898 | No | SRR5224139 | - | SRX2533320 |
| BRF1 | hESC H9 | 67311194 | 16129317 | No | SRR5224136 | - | SRX2533317 |
| GTF3C1 | HeLa | 88466227 | 9241855 | No | SRR036662 | - | SRX017005 |
| GTF3C1 | HeLa | 88466227 | 14248117 | No | SRR039064 | - | SRX018608 |
| GTF3C1 | HeLa | 88466227 | 11753968 | No | SRR039065 | - | SRX018608 |
| GTF3C1 | HeLa | 88466227 | 22169137 | No | SRR502068 | - | SRX150422 |
| GTF3C1 | HeLa | 88466227 | 8933197 | No | SRR036663 | - | SRX017005 |
| GTF3C1 | HeLa | 88466227 | 22119953 | No | SRR502069 | - | SRX150422 |
| GTF3C1 | hESC H9 | 54391172 | 9501628 | No | SRR5224141 | - | SRX2533322 |
| GTF3C1 | hESC H9 | 54391172 | 9504916 | No | SRR5224140 | - | SRX2533321 |
| GTF3C1 | hESC H9 | 54391172 | 7230666 | No | SRR5224142 | - | SRX2533323 |
| GTF3C1 | hESC H9 | 54391172 | 7070482 | No | SRR5224143 | - | SRX2533323 |
| GTF3C1 | hESC H9 | 54391172 | 10703233 | No | SRR5224144 | - | SRX2533324 |
| GTF3C1 | hESC H9 | 54391172 | 10380247 | No | SRR5224145 | - | SRX2533324 |
| GTF3C2 | IMR-90 | 63693990 | 17387155 | No | SRR7875612 | - | SRX4714471 |
| GTF3C2 | IMR-90 | 63693990 | 46306835 | No | SRR7875614 | - | SRX4714473 |
| GTF3C2 | T-47D | 346434711 | 120642641 | No | SRR7875596 | - | SRX4714455 |
| GTF3C2 | T-47D | 346434711 | 48147085 | No | SRR7875621 | - | SRX4714480 |
| GTF3C2 | T-47D | 346434711 | 57175466 | No | SRR9886581 | - | SRX6639037 |
| GTF3C2 | T-47D | 346434711 | 56658746 | No | SRR9886582 | - | SRX6639038 |
| GTF3C2 | T-47D | 346434711 | 63810773 | No | SRR7875592 | - | SRX4714451 |
| GTF3C2 | T98G | 225527505 | 84498530 | No | SRR7875608 | - | SRX4714467 |
| GTF3C2 | T98G | 225527505 | 70348441 | No | SRR7875610 | - | SRX4714469 |
| GTF3C2 | T98G | 225527505 | 70680534 | No | SRR7875623 | - | SRX4714482 |
| GTF3C2 | hESC H9 | 64899391 | 64899391 | No | SRR17771681 | - | SRX13933973 |
| GTF3C5 | IMR-5 | 20339183 | 10114276 | No | SRR3207449 | - | SRX1616411 |
| GTF3C5 | IMR-5 | 20339183 | 10224907 | No | SRR3207450 | - | SRX1616412 |
| IGG | A549 | 157264642 | 80509334 | Yes | ENCFF002EJF | ENCFF002EJD |  |
| IGG | A549 | 157264642 | 76755308 | Yes | ENCFF002EJI | ENCFF002EJB |  |
| IGG | HepG2 | 151907668 | 72117212 | Yes | ENCFF002EIA | ENCFF002EID |  |
| IGG | HepG2 | 151907668 | 79790456 | Yes | ENCFF002EIB | ENCFF002EIC |  |
| IGG | K562 | 105419078 | 39738737 | Yes | ENCFF002EFY | ENCFF002DUC |  |
| IGG | K562 | 105419078 | 65680341 | Yes | ENCFF002EGI | ENCFF002EGA |  |
| IGG | -MCF 7.00 | 314218130 | 90972566 | Yes | ENCFF002EHD | ENCFF002DTX |  |
| IGG | -MCF 7.00 | 314218130 | 59635093 | Yes | ENCFF002ELW | ENCFF002ELX |  |
| IGG | -MCF 7.00 | 314218130 | 73676920 | Yes | ENCFF002DUF | ENCFF002EGX |  |
| IGG | -MCF 7.00 | 314218130 | 89933551 | Yes | ENCFF002EHE | ENCFF002EHF |  |
| IGG | THP-1 | 69948627 | 69948627 | Yes | SRR13268105_1 | SRR13268105_2 |  |
| POLR1B | HEK293T | 52952799 | 11860696 | No | SRR087747 | - |  |
| POLR1A | K562 | 52952799 | 41092103 | No | ERR2109141 | - |  |
| POLR2A | A549 | 69668888 | 36992512 | No | ENCFF000RMM | - |  |
| POLR2A | A549 | 69668888 | 32676376 | No | ENCFF000RMO | - |  |
| POLR2A | GM12878 | 139912938 | 5714798 | No | ENCFF000WBQ | - |  |
| POLR2A | GM12878 | 139912938 | 6004938 | No | ENCFF000WBS | - |  |
| POLR2A | GM12878 | 139912938 | 5368875 | No | ENCFF000WBT | - |  |
| POLR2A | GM12878 | 139912938 | 6972753 | No | ENCFF000WBV | - |  |
| POLR2A | GM12878 | 139912938 | 5672214 | No | ENCFF000WBW | - |  |
| POLR2A | GM12878 | 139912938 | 6336821 | No | ENCFF000WBX | - |  |
| POLR2A | GM12878 | 139912938 | 14795146 | No | ENCFF000WCC | - |  |
| POLR2A | GM12878 | 139912938 | 16157397 | No | ENCFF000RPI | - |  |
| POLR2A | GM12878 | 139912938 | 11017478 | No | ENCFF000RPL | - |  |
| POLR2A | GM12878 | 139912938 | 31454718 | No | ENCFF000OBB | - |  |
| POLR2A | GM12878 | 139912938 | 30417800 | No | ENCFF000OBE | - |  |
| POLR2A | GM12891 | 109711035 | 19824077 | No | ENCFF000OIW | - |  |
| POLR2A | GM12891 | 109711035 | 21873581 | No | ENCFF000OIZ | - |  |

|  |  |  |  |  |  |  |
| --- | --- | --- | --- | --- | --- | --- |
| POLR2A | GM12891 | 109711035 | 14182328 | No | ENCFF000WIL | - |
| POLR2A | GM12891 | 109711035 | 16011135 | No | ENCFF000WIM | - |
| POLR2A | GM12891 | 109711035 | 16430525 | No | ENCFF000WIO | - |
| POLR2A | GM12891 | 109711035 | 6640065 | No | ENCFF000WIW | - |
| POLR2A | GM12891 | 109711035 | 7429710 | No | ENCFF000WJB | - |
| POLR2A | GM12891 | 109711035 | 7319614 | No | ENCFF000WJC | - |
| POLR2A | GM12892 | 80193877 | 6320354 | No | ENCFF000WJQ | - |
| POLR2A | GM12892 | 80193877 | 8561445 | No | ENCFF000WJR | - |
| POLR2A | GM12892 | 80193877 | 10752673 | No | ENCFF000WJY | - |
| POLR2A | GM12892 | 80193877 | 21165664 | No | ENCFF000OLH | - |
| POLR2A | GM12892 | 80193877 | 33393741 | No | ENCFF000OLJ | - |
| POLR2A | GM18505 | 37137423 | 6248887 | No | ENCFF000WLY | - |
| POLR2A | GM18505 | 37137423 | 6316977 | No | ENCFF000WMB | - |
| POLR2A | GM18505 | 37137423 | 10699173 | No | ENCFF000WMC | - |
| POLR2A | GM18505 | 37137423 | 13872386 | No | ENCFF000WML | - |
| POLR2A | GM18951 | 24180945 | 4969650 | No | ENCFF000WOB | - |
| POLR2A | GM18951 | 24180945 | 6977178 | No | ENCFF000WOC | - |
| POLR2A | GM18951 | 24180945 | 3664853 | No | ENCFF000WOF | - |
| POLR2A | GM18951 | 24180945 | 8569264 | No | ENCFF000WOL | - |
| POLR2A | GM19099 | 28056135 | 6990880 | No | ENCFF000WPB | - |
| POLR2A | GM19099 | 28056135 | 5623514 | No | ENCFF000WPD | - |
| POLR2A | GM19099 | 28056135 | 15441741 | No | ENCFF000WPL | - |
| POLR2A | GM19193 | 25626801 | 4370301 | No | ENCFF000WQF | - |
| POLR2A | GM19193 | 25626801 | 4873260 | No | ENCFF000WQJ | - |
| POLR2A | GM19193 | 25626801 | 7578532 | No | ENCFF000WQM | - |
| POLR2A | GM19193 | 25626801 | 8804708 | No | ENCFF000WRO | - |
| POLR2A | H1 | 73956989 | 33104798 | No | ENCFF000ORP | - |
| POLR2A | H1 | 73956989 | 23331771 | No | ENCFF000ORR | - |
| POLR2A | H1 | 73956989 | 17520420 | No | ENCFF000RSM | - |
| POLR2A | H54 | 36054194 | 20037709 | No | ENCFF000RNL | - |

|  |  |  |  |  |  |  |
| --- | --- | --- | --- | --- | --- | --- |
| POLR2A | H54 | 36054194 | 16016485 | No | ENCFF000RNQ | - |
| POLR2A | HeLa-S3 | 87340517 | 5482275 | No | ENCFF000RTM | - |
| POLR2A | HeLa-S3 | 87340517 | 24693639 | No | ENCFF000PEV | - |
| POLR2A | HeLa-S3 | 87340517 | 29265379 | No | ENCFF000PFC | - |
| POLR2A | HeLa-S3 | 87340517 | 7646887 | No | ENCFF000XJP | - |
| POLR2A | HeLa-S3 | 87340517 | 10259928 | No | ENCFF000XJS | - |
| POLR2A | HeLa-S3 | 87340517 | 9992409 | No | ENCFF000XKD | - |
| POLR2A | HepG2 | 105583880 | 17379821 | No | ENCFF000PNM | - |
| POLR2A | HepG2 | 105583880 | 25447057 | No | ENCFF000PNT | - |
| POLR2A | HepG2 | 105583880 | 18964839 | No | ENCFF000XWJ | - |
| POLR2A | HepG2 | 105583880 | 23246385 | No | ENCFF000XWR | - |
| POLR2A | HepG2 | 105583880 | 5204101 | No | ENCFF000RUR | - |
| POLR2A | HepG2 | 105583880 | 7043434 | No | ENCFF000RUT | - |
| POLR2A | HepG2 | 105583880 | 8298243 | No | ENCFF000RUY | - |
| POLR2A | IMR-90 | 54368055 | 27743747 | No | ENCFF000YDX | - |
| POLR2A | IMR-90 | 54368055 | 26624308 | No | ENCFF000YDZ | - |
| POLR2A | K562 | 327866848 | 11304945 | No | ENCFF000YXN | - |
| POLR2A | K562 | 327866848 | 10635844 | No | ENCFF000YXO | - |
| POLR2A | K562 | 327866848 | 34224229 | No | ENCFF000QDX | - |
| POLR2A | K562 | 327866848 | 32195895 | No | ENCFF000QDY | - |
| POLR2A | K562 | 327866848 | 48941971 | Yes | ENCFF959SCN | ENCFF545PSD |
| POLR2A | K562 | 327866848 | 51075620 | Yes | ENCFF272EEG | ENCFF531FHF |
| POLR2A | K562 | 327866848 | 8984193 | No | ENCFF000YXG | - |
| POLR2A | K562 | 327866848 | 9565532 | No | ENCFF000YXH | - |
| POLR2A | K562 | 327866848 | 255 | No | ENCFF000RWV | - |
| POLR2A | K562 | 327866848 | 17291350 | No | ENCFF000RXB | - |
| POLR2A | K562 | 327866848 | 58713927 | Yes | ENCFF839LPL | ENCFF698ICA |
| POLR2A | K562 | 327866848 | 44933087 | Yes | ENCFF920SBM | ENCFF400NVU |
| POLR2A | -MCF 7.00 | 200053553 | 36340241 | No | ENCFF000SBV | - |
| POLR2A | -MCF 7.00 | 200053553 | 36311353 | No | ENCFF000SBY | - |

|  |  |  |  |  |  |  |
| --- | --- | --- | --- | --- | --- | --- |
| POLR2A | -MCF 7.00 | 200053553 | 34552609 | No | ENCFF000SBO | - |
| POLR2A | -MCF 7.00 | 200053553 | 35572561 | No | ENCFF000SBP | - |
| POLR2A | -MCF 7.00 | 200053553 | 29175131 | No | ENCFF000SBI | - |
| POLR2A | -MCF 7.00 | 200053553 | 28101658 | No | ENCFF000SBK | - |
| POLR2A | Raji | 22497988 | 7399747 | No | ENCFF000ZPU | - |
| POLR2A | Raji | 22497988 | 15098241 | No | ENCFF000ZQE | - |
| POLR2A | adrenal gland | 225552181 | 93365718 | Yes | ENCFF411THN | ENCFF180UGD |
| POLR2A | adrenal gland | 225552181 | 132186463 | Yes | ENCFF933ONW | ENCFF464OZG |
| POLR2A | body of pancreas | 446409687 | 123977836 | Yes | ENCFF943DRV | ENCFF195OVM |
| POLR2A | body of pancreas | 446409687 | 158988031 | Yes | ENCFF703TVC | ENCFF826QIL |
| POLR2A | body of pancreas | 446409687 | 67221458 | Yes | ENCFF176BWH | ENCFF019HPW |
| POLR2A | body of pancreas | 446409687 | 96222362 | Yes | ENCFF506ONU | ENCFF113JGC |
| POLR2A | breast epithelium | 130525366 | 54994093 | Yes | ENCFF074WTD | ENCFF784DME |
| POLR2A | breast epithelium | 130525366 | 75531273 | Yes | ENCFF177KGH | ENCFF225AAI |
| POLR2A | endothelial cell of umbilical vein | 82191493 | 136780 | No | ENCFF000RVR | - |
| POLR2A | endothelial cell of umbilical vein | 82191493 | 403145 | No | ENCFF000RVW | - |
| POLR2A | endothelial cell of umbilical vein | 82191493 | 10950397 | No | ENCFF000YBD | - |
| POLR2A | endothelial cell of umbilical vein | 82191493 | 11042185 | No | ENCFF000YBR | - |
| POLR2A | endothelial cell of umbilical vein | 82191493 | 9553993 | No | ENCFF000YCV | - |
| POLR2A | endothelial cell of umbilical vein | 82191493 | 14988670 | No | ENCFF000PVK | - |
| POLR2A | endothelial cell of umbilical vein | 82191493 | 35116323 | No | ENCFF000PVP | - |
| POLR2A | erythroblast | 64566593 | 33581060 | No | ENCFF000ZPK | - |
| POLR2A | erythroblast | 64566593 | 30985533 | No | ENCFF000ZPN | - |
| POLR2A | esophagus muscularis mucosa | 71140434 | 71140434 | Yes | ENCFF438UOR | ENCFF478FFG |
| POLR2A | esophagus squamous epithelium | 354133036 | 100268058 | Yes | ENCFF813TWG | ENCFF141YYX |
| POLR2A | esophagus squamous epithelium | 354133036 | 81931897 | Yes | ENCFF547OPO | ENCFF996YRU |
| POLR2A | esophagus squamous epithelium | 354133036 | 84185075 | Yes | ENCFF356QEU | ENCFF759FBS |
| POLR2A | esophagus squamous epithelium | 354133036 | 87748006 | Yes | ENCFF096WFG | ENCFF550IEW |
| POLR2A | gastrocnemius medialis | 172644056 | 80198288 | Yes | ENCFF956QIS | ENCFF300RLH |
| POLR2A | gastrocnemius medialis | 172644056 | 92445768 | Yes | ENCFF570FCV | ENCFF713HTL |

|  |  |  |  |  |  |  |
| --- | --- | --- | --- | --- | --- | --- |
| POLR2A | gastroesophageal sphincter | 334803278 | 41713303 | Yes | ENCFF763EWZ | ENCFF828DEK |
| POLR2A | gastroesophageal sphincter | 334803278 | 41819349 | Yes | ENCFF331BQR | ENCFF082IYB |
| POLR2A | gastroesophageal sphincter | 334803278 | 90069276 | Yes | ENCFF545SYX | ENCFF972WGI |
| POLR2A | gastroesophageal sphincter | 334803278 | 58104624 | Yes | ENCFF026VCA | ENCFF357YNX |
| POLR2A | gastroesophageal sphincter | 334803278 | 103096726 | Yes | ENCFF336GML | ENCFF163KIH |
| POLR2A | heart left ventricle | 173047716 | 92559249 | Yes | ENCFF791UGB | ENCFF254TVF |
| POLR2A | heart left ventricle | 173047716 | 80488467 | Yes | ENCFF999VOH | ENCFF435KUB |
| POLR2A | lower leg skin | 463795080 | 92054194 | Yes | ENCFF068NIQ | ENCFF844DFN |
| POLR2A | lower leg skin | 463795080 | 100700344 | Yes | ENCFF599ZRC | ENCFF229BOO |
| POLR2A | lower leg skin | 463795080 | 76436042 | Yes | ENCFF641KSE | ENCFF246RFD |
| POLR2A | lower leg skin | 463795080 | 127308342 | Yes | ENCFF346MYJ | ENCFF223FTQ |
| POLR2A | lower leg skin | 463795080 | 67296158 | Yes | ENCFF182BFU | ENCFF967GHE |
| POLR2A | ovary | 102416246 | 102416246 | Yes | ENCFF384MEM | ENCFF322TJU |
| POLR2A | prostate gland | 174920444 | 79057976 | Yes | ENCFF883DYX | ENCFF516YRH |
| POLR2A | prostate gland | 174920444 | 95862468 | Yes | ENCFF278XHE | ENCFF869NYZ |
| POLR2A | right lobe of liver | 69246367 | 69246367 | Yes | ENCFF174GER | ENCFF071CTT |
| POLR2A | sigmoid colon | 301285138 | 91865667 | Yes | ENCFF843LGK | ENCFF353IIG |
| POLR2A | sigmoid colon | 301285138 | 73581255 | Yes | ENCFF943KNW | ENCFF509ERN |
| POLR2A | sigmoid colon | 301285138 | 92248579 | Yes | ENCFF015MWE | ENCFF405OKO |
| POLR2A | sigmoid colon | 301285138 | 43589637 | Yes | ENCFF112BSI | ENCFF160QJZ |
| POLR2A | spleen | 391884237 | 119822087 | Yes | ENCFF073GNS | ENCFF975DHQ |
| POLR2A | spleen | 391884237 | 121258426 | Yes | ENCFF982FBQ | ENCFF837IQA |
| POLR2A | spleen | 391884237 | 73443057 | Yes | ENCFF510UIJ | ENCFF606RQN |
| POLR2A | spleen | 391884237 | 77360667 | Yes | ENCFF894LNI | ENCFF019BTC |
| POLR2A | stomach | 336112177 | 72234850 | Yes | ENCFF699OSQ | ENCFF014DFT |
| POLR2A | stomach | 336112177 | 60903238 | Yes | ENCFF800HLE | ENCFF700SNB |
| POLR2A | stomach | 336112177 | 66182759 | Yes | ENCFF073GTF | ENCFF554PAM |
| POLR2A | stomach | 336112177 | 43005346 | Yes | ENCFF090TRF | ENCFF014FRE |
| POLR2A | stomach | 336112177 | 93785984 | Yes | ENCFF800XBW | ENCFF233QXS |
| POLR2A | suprapubic skin | 196438792 | 110803932 | Yes | ENCFF310UBW | ENCFF834BMO |
| POLR2A | suprapubic skin | 196438792 | 85634860 | Yes | ENCFF706ERC | ENCFF797OQU |  |
| POLR2A | testis | 83265085 | 83265085 | Yes | ENCFF788VTV | ENCFF768RJW |  |
| POLR2A | thyroid gland | 199544925 | 118490417 | Yes | ENCFF536HLB | ENCFF669LRI |  |
| POLR2A | thyroid gland | 199544925 | 81054508 | Yes | ENCFF757IXD | ENCFF643EOI |  |
| POLR2A | tibial nerve | 158782236 | 74942111 | Yes | ENCFF556MUX | ENCFF672XQY |  |
| POLR2A | tibial nerve | 158782236 | 83840125 | Yes | ENCFF732EAG | ENCFF807YBJ |  |
| POLR2A | transverse colon | 261042982 | 114903528 | Yes | ENCFF445QWT | ENCFF384DGC |  |
| POLR2A | transverse colon | 261042982 | 71998308 | Yes | ENCFF945PBZ | ENCFF252EVO |  |
| POLR2A | transverse colon | 261042982 | 36770063 | Yes | ENCFF272HAB | ENCFF763GOA |  |
| POLR2A | transverse colon | 261042982 | 37371083 | Yes | ENCFF466DDM | ENCFF768VKW |  |
| POLR2A | upper lobe of left lung | 316893809 | 64749762 | Yes | ENCFF715ESG | ENCFF740EMX |  |
| POLR2A | upper lobe of left lung | 316893809 | 90434595 | Yes | ENCFF241CFU | ENCFF962IZX |  |
| POLR2A | upper lobe of left lung | 316893809 | 75103347 | Yes | ENCFF063DDH | ENCFF955ITN |  |
| POLR2A | upper lobe of left lung | 316893809 | 86606105 | Yes | ENCFF012WDS | ENCFF347JRV |  |
| POLR2A | uterus | 173973132 | 95600590 | Yes | ENCFF500XDC | ENCFF042CMS |  |
| POLR2A | uterus | 173973132 | 78372542 | Yes | ENCFF973HMK | ENCFF534XBR |  |
| POLR2A | vagina | 164935104 | 65839982 | Yes | ENCFF166CFL | ENCFF115IMK |  |
| POLR2A | vagina | 164935104 | 99095122 | Yes | ENCFF389ZTP | ENCFF788KCN |  |
| POLR3A | 293 | 417875684 | 90451568 | Yes | SRR7717420_1 | SRR7717420_2 | SRX4574092 |
| POLR3A | 293 | 417875684 | 110822913 | Yes | SRR7717421_1 | SRR7717421_2 | SRX4574093 |
| POLR3A | 293 | 417875684 | 105424077 | Yes | SRR7717419_1 | SRR7717419_2 | SRX4574091 |
| POLR3A | 293 | 417875684 | 111177126 | Yes | SRR7717422_1 | SRR7717422_2 | SRX4574094 |
| POLR3A | THP-1 | 158290549 | 46681820 | Yes | SRR13268085_1 | SRR13268085_2 | SRX9698427 |
| POLR3A | THP-1 | 158290549 | 18991432 | Yes | SRR13268086_1 | SRR13268086_2 | SRX9698427 |
| POLR3A | THP-1 | 158290549 | 25820847 | Yes | SRR13268088_1 | SRR13268088_2 | SRX9698428 |
| POLR3A | THP-1 | 158290549 | 66796450 | Yes | SRR13268087_1 | SRR13268087_2 | SRX9698428 |
| POLR3B | THP-1 | 345917166 | 101409231 | Yes | SRR13268090_1 | SRR13268090_2 | SRX9698430 |
| POLR3B | THP-1 | 345917166 | 80162623 | Yes | SRR13268089_1 | SRR13268089_2 | SRX9698429 |
| POLR3B | THP-1 | 345917166 | 85374313 | Yes | SRR13268107_1 | SRR13268107_2 | SRX9698447 |
| POLR3B | THP-1 | 345917166 | 78970999 | Yes | SRR13268106_1 | SRR13268106_2 | SRX9698446 |
| POLR3C | 293 | 337277136 | 85834537 | No | SRR10990458 | - | SRX5803016 |
| POLR3C | 293 | 337277136 | 81293731 | No | SRR10990461 | - | SRX5803019 |
| POLR3C | 293 | 337277136 | 80407568 | No | SRR10990459 | - | SRX5803017 |
| POLR3C | 293 | 337277136 | 89741300 | No | SRR10990460 | - | SRX5803018 |
| POLR3C | THP-1 | 277552948 | 127836523 | Yes | SRR13268093_1 | SRR13268093_2 | SRX9698433 |
| POLR3C | THP-1 | 277552948 | 149716425 | Yes | SRR13268094_1 | SRR13268094_2 | SRX9698434 |
| POLR3D | IMR-90 | 2318624371 | 56951831 | No | SRR896108 | - | SRX301446 |
| POLR3D | IMR-90 | 2318624371 | 194380026 | No | SRR2633803 | - | SRX1330892 |
| POLR3D | IMR-90 | 2318624371 | 217695630 | No | SRR2633801 | - | SRX1330890 |
| POLR3D | IMR-90 | 2318624371 | 190762843 | No | SRR2633802 | - | SRX1330891 |
| POLR3D | IMR-90 | 2318624371 | 226110635 | No | SRR2633807 | - | SRX1330896 |
| POLR3D | IMR-90 | 2318624371 | 227517813 | No | SRR2633808 | - | SRX1330897 |
| POLR3D | IMR-90 | 2318624371 | 214884906 | No | SRR2633805 | - | SRX1330894 |
| POLR3D | IMR-90 | 2318624371 | 211344680 | No | SRR2633804 | - | SRX1330893 |
| POLR3D | IMR-90 | 2318624371 | 186713518 | No | SRR2633817 | - | SRX1330906 |
| POLR3D | IMR-90 | 2318624371 | 199971505 | No | SRR2633806 | - | SRX1330895 |
| POLR3D | IMR-90 | 2318624371 | 203989351 | No | SRR2633819 | - | SRX1330908 |
| POLR3D | IMR-90 | 2318624371 | 188301633 | No | SRR2633818 | - | SRX1330907 |
| POLR3D | THP-1 | 573670006 | 55606520 | Yes | SRR5359608_1 | SRR5359608_2 | SRX2655447 |
| POLR3D | THP-1 | 573670006 | 63564974 | Yes | SRR5359609_1 | SRR5359609_2 | SRX2655448 |
| POLR3D | THP-1 | 573670006 | 79328582 | Yes | SRR5359611_1 | SRR5359611_2 | SRX2655450 |
| POLR3D | THP-1 | 573670006 | 48030948 | Yes | SRR5359610_1 | SRR5359610_2 | SRX2655449 |
| POLR3D | THP-1 | 573670006 | 68090581 | Yes | SRR13268099_1 | SRR13268099_2 | SRX9698439 |
| POLR3D | THP-1 | 573670006 | 88168648 | Yes | SRR13268100_1 | SRR13268100_2 | SRX9698440 |
| POLR3D | THP-1 | 573670006 | 80327410 | Yes | SRR13268115_1 | SRR13268115_2 | SRX9698455 |
| POLR3D | THP-1 | 573670006 | 90552343 | Yes | SRR13268114_1 | SRR13268114_2 | SRX9698454 |
| POLR3E | THP-1 | 229954889 | 131119489 | Yes | SRR13268102_1 | SRR13268102_2 | SRX9698442 |
| POLR3E | THP-1 | 229954889 | 98835400 | Yes | SRR13268101_1 | SRR13268101_2 | SRX9698441 |
| POLR3G | IMR-90 | 55724283 | 55724283 | No | SRR896109 | - | SRX301447 |
| POLR3G | THP-1 | 339182841 | 44659853 | Yes | SRR13268095_1 | SRR13268095_2 | SRX9698435 |
| POLR3G | THP-1 | 339182841 | 62710285 | Yes | SRR13268096_1 | SRR13268096_2 | SRX9698436 |
| POLR3G | THP-1 | 339182841 | 48507178 | Yes | SRR13268110_1 | SRR13268110_2 | SRX9698450 |
| POLR3G | THP-1 | 339182841 | 55771545 | Yes | SRR13268111_1 | SRR13268111_2 | SRX9698451 |
| POLR3G | THP-1 | 339182841 | 56293998 | Yes | SRR13268116_1 | SRR13268116_2 | SRX9698456 |
| POLR3G | THP-1 | 339182841 | 71239982 | Yes | SRR13268117_1 | SRR13268117_2 | SRX9698457 |
| POLR3GL | IMR-90 | 54875000 | 54875000 | No | SRR896110 | - | SRX301448 |
| POLR3GL | THP-1 | 466478517 | 73676314 | Yes | SRR13268097_1 | SRR13268097_2 | SRX9698437 |
| POLR3GL | THP-1 | 466478517 | 80569182 | Yes | SRR13268098_1 | SRR13268098_2 | SRX9698438 |
| POLR3GL | THP-1 | 466478517 | 78956199 | Yes | SRR13268113_1 | SRR13268113_2 | SRX9698453 |
| POLR3GL | THP-1 | 466478517 | 70879453 | Yes | SRR13268112_1 | SRR13268112_2 | SRX9698452 |
| POLR3GL | THP-1 | 466478517 | 84928225 | Yes | SRR13268119_1 | SRR13268119_2 | SRX9698459 |
| POLR3GL | THP-1 | 466478517 | 77469144 | Yes | SRR13268118_1 | SRR13268118_2 | SRX9698458 |
| SNAPC1 | IMR-90 | 29017923 | 29017923 | No | SRR501634 | - | SRX150108 |
| SNAPC2 | IMR-90 | 30177585 | 4840610 | No | SRR035559 | - | SRX016559 |
| SNAPC2 | IMR-90 | 30177585 | 11508985 | No | SRR501633 | - | SRX150107 |
| SNAPC2 | IMR-90 | 30177585 | 9839394 | No | SRR501632 | - | SRX150106 |
| SNAPC2 | IMR-90 | 30177585 | 3988596 | No | SRR035558 | - | SRX016559 |
| SNAPC4 | HCT 116 | 84109060 | 43836738 | No | SRR11969088 | - | SRX8512868 |
| SNAPC4 | HCT 116 | 84109060 | 40272322 | No | SRR11969089 | - | SRX8512869 |
| SNAPC4 | Hep G2 | 79436906 | 40342209 | No | SRR14104370 | - | SRX10475809 |
| SNAPC4 | Hep G2 | 79436906 | 39094697 | No | SRR14104371 | - | SRX10475810 |
| SNAPC4 | LoVo | 22451100 | 22451100 | No | SRR1002575 | - | SRX360040 |
| SNAPC4 | MCF 10A | 29935326 | 29935326 | No | SRR488764 | - | SRX143300 |
| SNAPC5 | IMR-90 | 31799535 | 31799535 | No | SRR501635 | - | SRX150109 |
| SNAPC5 | K-562 | 190556472 | 72086357 | Yes | SRR14636379_1 | SRR14636379_2 | SRX10976129 |
| SNAPC5 | K-562 | 190556472 | 118470115 | Yes | SRR14636380_1 | SRR14636380_2 | SRX10976130 |
| TBP | 293 | 96140178 | 49730483 | Yes | SRR14879798_1 | SRR14879798_2 | SRX11196941 |
| TBP | 293 | 96140178 | 46409695 | Yes | SRR14879799_1 | SRR14879799_2 | SRX11196942 |
| TBP | Bronchial tracheal epithelial cells | 84987491 | 32114691 | No | SRR7497981 | - | SRX4367422 |
| TBP | Bronchial tracheal epithelial cells | 84987491 | 17651376 | No | SRR7497980 | - | SRX4367421 |
| TBP | Bronchial tracheal epithelial cells | 84987491 | 9979248 | No | SRR7498000 | - | SRX4367441 |
| TBP | Bronchial tracheal epithelial cells | 84987491 | 8866343 | No | SRR7497990 | - | SRX4367431 |
| TBP | Bronchial tracheal epithelial cells | 84987491 | 10411362 | No | SRR7498001 | - | SRX4367442 |
| TBP | Bronchial tracheal epithelial cells | 84987491 | 5964471 | No | SRR7497991 | - | SRX4367432 |
| TBP | Foreskin | 320150768 | 89680353 | Yes | SRR16288497_1 | SRR16288497_2 | SRX12567770 |
| TBP | Foreskin | 320150768 | 88592298 | Yes | SRR16288498_1 | SRR16288498_2 | SRX12567770 |
| TBP | Foreskin | 320150768 | 76349960 | Yes | SRR16288509_1 | SRR16288509_2 | SRX12567778 |
| TBP | Foreskin | 320150768 | 65528157 | Yes | SRR16288510_1 | SRR16288510_2 | SRX12567779 |
| TBP | GM12878 | 44009147 | 22685432 | No | SRR501925 | - | SRX150357 |
| TBP | GM12878 | 44009147 | 21323715 | No | SRR501926 | - | SRX150357 |
| TBP | HeLa | 38430877 | 20812978 | No | SRR502618 | - | SRX150685 |
| TBP | HeLa | 38430877 | 17617899 | No | SRR502619 | - | SRX150685 |
| TBP | Hep G2 | 109256620 | 21329866 | No | SRR501933 | - | SRX150360 |
| TBP | Hep G2 | 109256620 | 22328235 | No | SRR501932 | - | SRX150360 |
| TBP | Hep G2 | 109256620 | 27266446 | No | SRR14103929 | - | SRX10475554 |
| TBP | Hep G2 | 109256620 | 38332073 | No | SRR14103928 | - | SRX10475553 |
| TBP | K-562 | 589418373 | 24624617 | No | SRR502379 | - | SRX150574 |
| TBP | K-562 | 589418373 | 22152184 | No | SRR502378 | - | SRX150574 |
| TBP | K-562 | 589418373 | 22943525 | No | SRR1185445 | - | SRX482859 |
| TBP | K-562 | 589418373 | 62555232 | Yes | SRR13049288_1 | SRR13049288_2 | SRX9498683 |
| TBP | K-562 | 589418373 | 46750034 | Yes | SRR13049291_1 | SRR13049291_2 | SRX9498686 |
| TBP | K-562 | 589418373 | 37541424 | Yes | SRR13049289_1 | SRR13049289_2 | SRX9498684 |
| TBP | K-562 | 589418373 | 71000320 | Yes | SRR13049268_1 | SRR13049268_2 | SRX9498730 |
| TBP | K-562 | 589418373 | 57376407 | Yes | SRR13049269_1 | SRR13049269_2 | SRX9498731 |
| TBP | K-562 | 589418373 | 78377485 | Yes | SRR13049270_1 | SRR13049270_2 | SRX9498732 |
| TBP | K-562 | 589418373 | 85899979 | Yes | SRR13049271_1 | SRR13049271_2 | SRX9498733 |
| TBP | K-562 | 589418373 | 80197166 | Yes | SRR13049290_1 | SRR13049290_2 | SRX9498685 |
| TBP | Mesenchymal stem cells | 75695050 | 41258851 | Yes | SRR12467672_1 | SRR12467672_2 | SRX8961847 |
| TBP | Mesenchymal stem cells | 75695050 | 34436199 | Yes | SRR12467683_1 | SRR12467683_2 | SRX8961858 |
| TBP | U2OS | 48109418 | 20720169 | No | SRR14370555 | - | SRX10722986 |
| TBP | U2OS | 48109418 | 27389249 | No | SRR14370556 | - | SRX10722987 |
| TBP | hESC H1 | 214136218 | 24793243 | No | SRR501988 | - | SRX150383 |
| TBP | hESC H1 | 214136218 | 29870381 | No | SRR501987 | - | SRX150383 |
| TBP | hESC H1 | 214136218 | 44738177 | No | SRR9185038 | - | SRX5957524 |
| TBP | hESC H1 | 214136218 | 39420588 | No | SRR9185039 | - | SRX5957525 |
| TBP | hESC H1 | 214136218 | 39404933 | No | SRR9185040 | - | SRX5957526 |
| TBP | hESC H1 | 214136218 | 35908896 | No | SRR9185041 | - | SRX5957527 |
| TBP | hESC H9 | 349866389 | 79210799 | Yes | SRR14802462_1 | SRR14802462_2 | SRX11135513 |
| TBP | hESC H9 | 349866389 | 95502636 | Yes | SRR14802464_1 | SRR14802464_2 | SRX11135515 |
| TBP | hESC H9 | 349866389 | 87162673 | Yes | SRR14802463_1 | SRR14802463_2 | SRX11135514 |
| TBP | hESC H9 | 349866389 | 87990281 | Yes | SRR14802465_1 | SRR14802465_2 | SRX11135516 |
| TBP | iPS cells | 257628585 | 17193378 | No | SRR11885039 | - | SRX8432576 |
| TBP | iPS cells | 257628585 | 16066035 | No | SRR11885040 | - | SRX8432576 |
| TBP | iPS cells | 257628585 | 16454130 | No | SRR11885042 | - | SRX8432577 |
| TBP | iPS cells | 257628585 | 17601779 | No | SRR11885041 | - | SRX8432577 |
| TBP | iPS cells | 257628585 | 15360117 | No | SRR11885044 | - | SRX8432578 |
| TBP | iPS cells | 257628585 | 16419122 | No | SRR11885043 | - | SRX8432578 |
| TBP | iPS cells | 257628585 | 16813490 | No | SRR11885045 | - | SRX8432579 |
| TBP | iPS cells | 257628585 | 15778470 | No | SRR11885046 | - | SRX8432579 |
| TBP | iPS cells | 257628585 | 17122751 | No | SRR11885047 | - | SRX8432580 |
| TBP | iPS cells | 257628585 | 16023459 | No | SRR11885048 | - | SRX8432580 |
| TBP | iPS cells | 257628585 | 16437172 | No | SRR11885049 | - | SRX8432584 |
| TBP | iPS cells | 257628585 | 15334502 | No | SRR11885050 | - | SRX8432584 |
| TBP | iPS cells | 257628585 | 16302030 | No | SRR11885051 | - | SRX8432585 |
| TBP | iPS cells | 257628585 | 15184997 | No | SRR11885052 | - | SRX8432585 |
| TBP | iPS cells | 257628585 | 15224782 | No | SRR11885053 | - | SRX8432586 |
| TBP | iPS cells | 257628585 | 14312371 | No | SRR11885054 | - | SRX8432586 |

**Supplementary Table 2.** PRO-seq data used in this study.

| Cell.type | paired | read1 | read2 | size_of_bam_file | fraction_of_unknown_specificity |
| --- | --- | --- | --- | --- | --- |
| CHM13,human hydatiform mole | No | SRR15035502 | - | 2.9G | 0.0728 |
| CHM13,human hydatiform mole | No | SRR15035503 | - | 3.5G | 0.0731 |
| Hela | Yes | SRR5799635_1 | SRR5799635_2 | 1.8G | 0.0665 |
| Hela-S3 | No | SRR15052413 | - | 1.1G | 0.0695 |
| Hela-S3 | No | SRR15052414 | - | 1.5G | 0.0673 |
| K562 | No | SRR1554311 | - | 11G | 0.0513 |
| K562 | No | SRR1554312 | - | 11G | 0.0513 |
| RPE1 | No | SRR15035500 | - | 1.2G | 0.0713 |
| RPE1 | No | SRR15035501 | - | 1.5G | 0.0717 |
| THP1 | Yes | SRR5359616_1 | SRR5359616_2 | 1.2G | 0.0433 |
| THP1 | Yes | SRR5359617_1 | SRR5359617_2 | 865M | 0.0529 |
| THP1+PMA72hr | Yes | SRR5359618_1 | SRR5359618_2 | 801M | 0.0498 |
| THP1+PMA72hr | Yes | SRR5359619_1 | SRR5359619_2 | 1.0G | 0.0462 |

**Supplementary Table 3.**
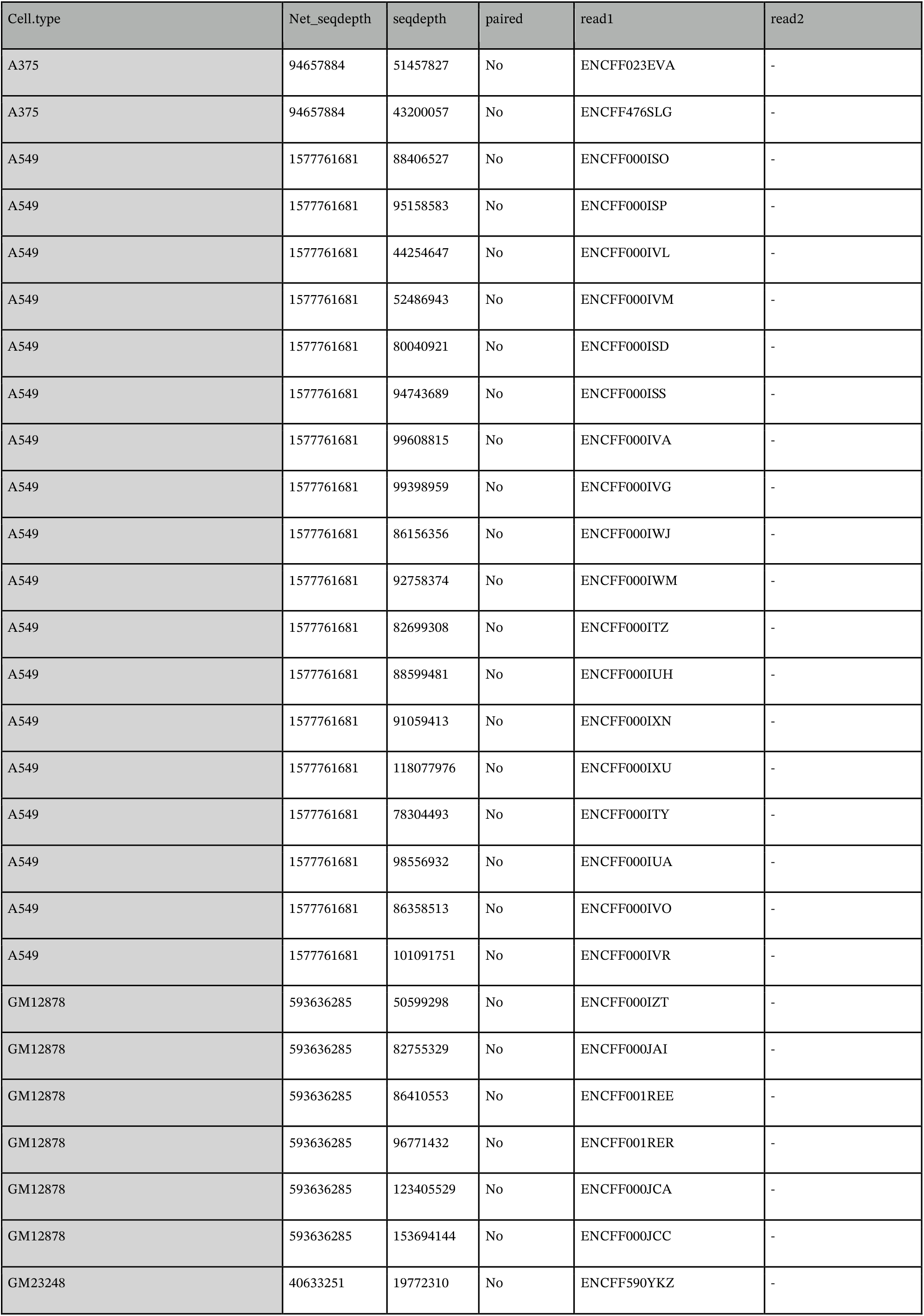
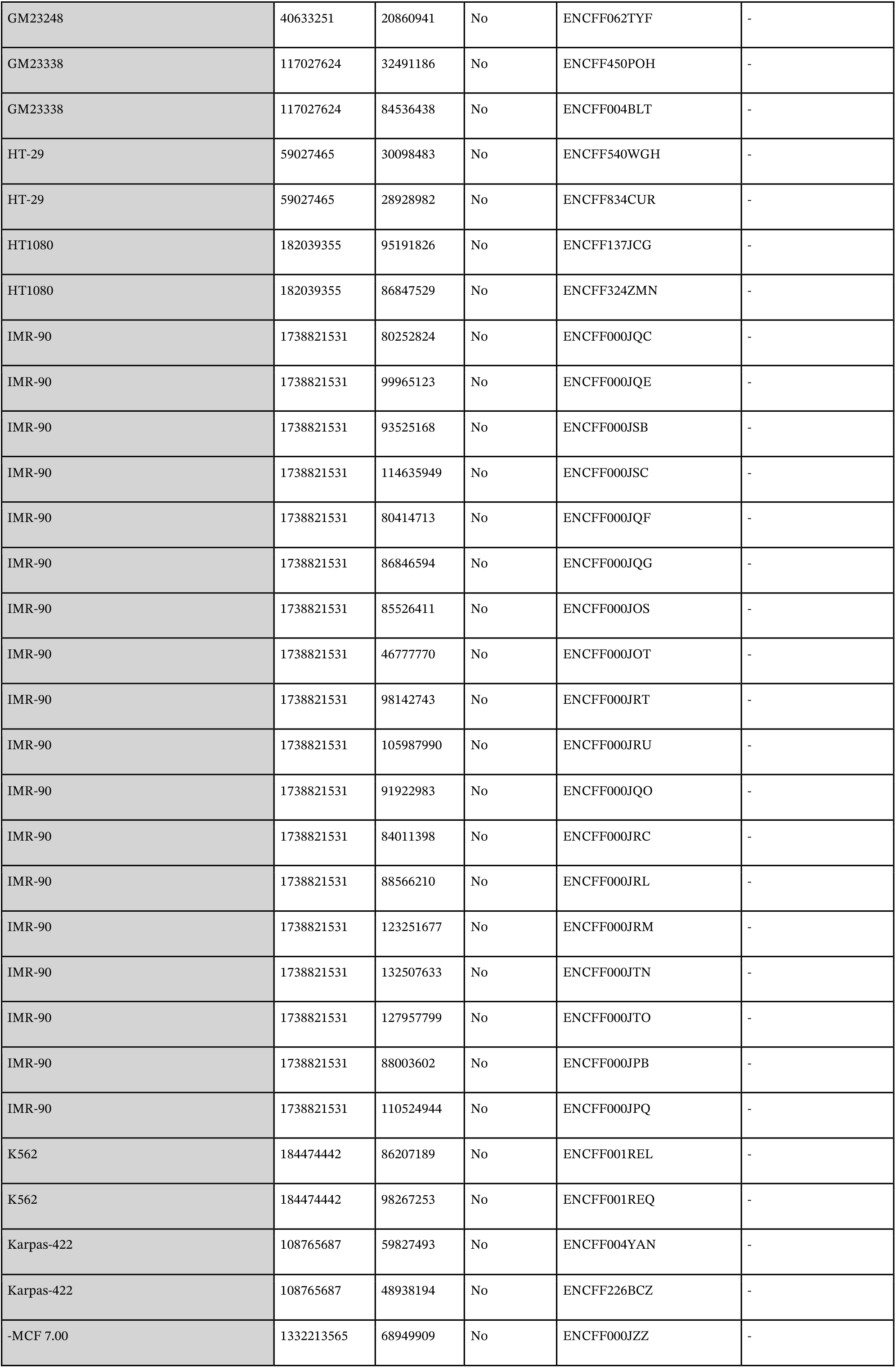
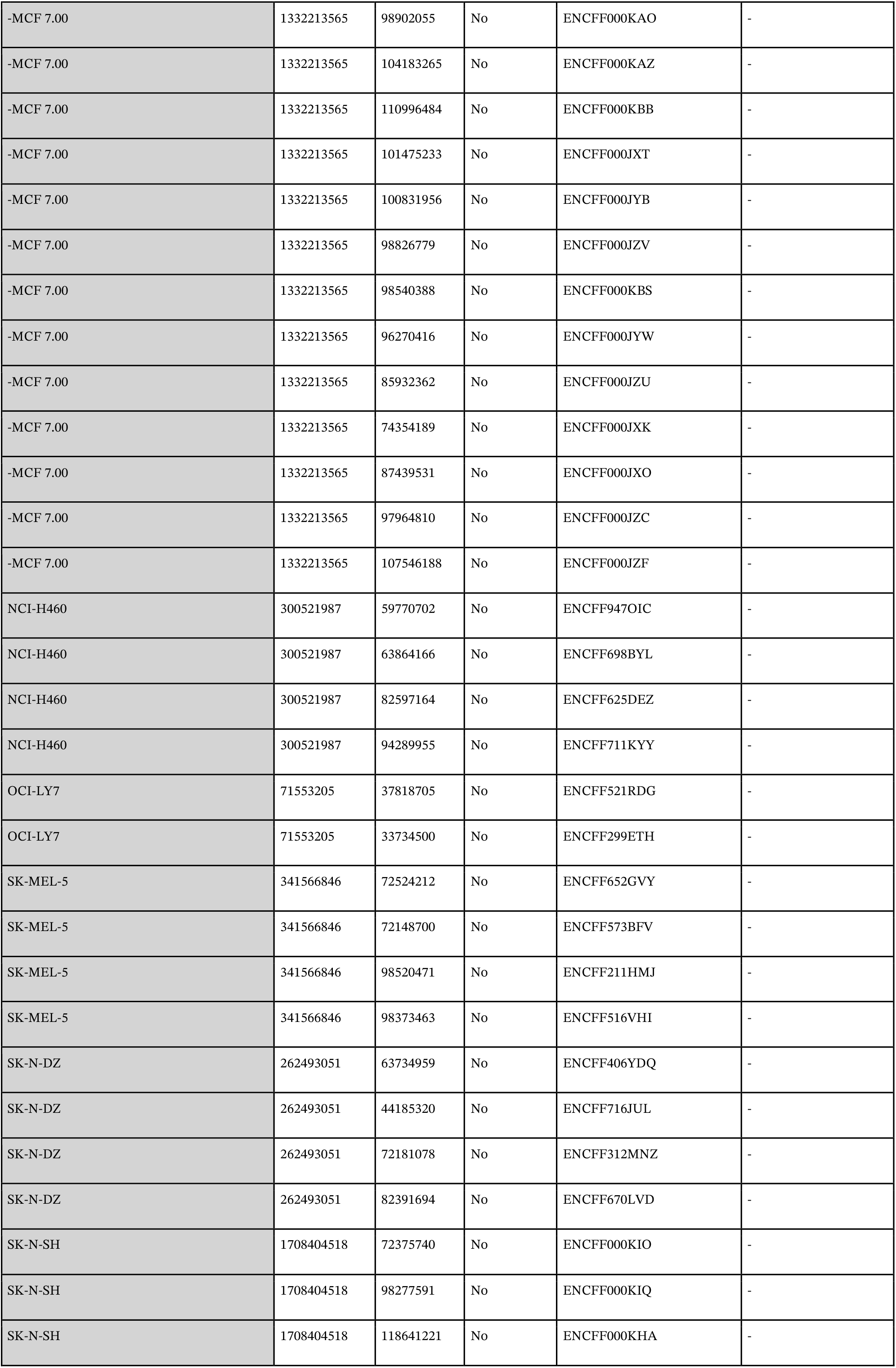
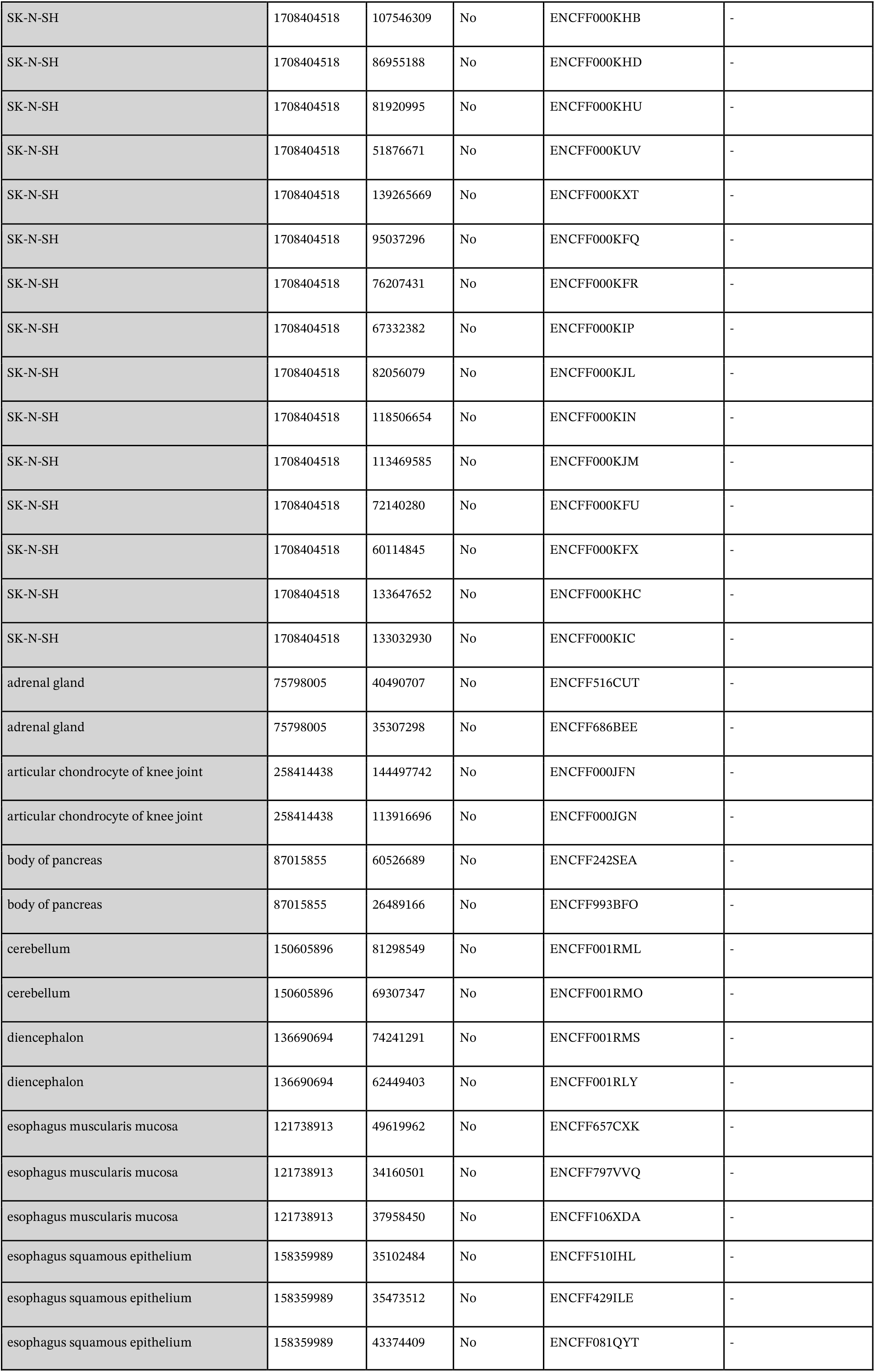
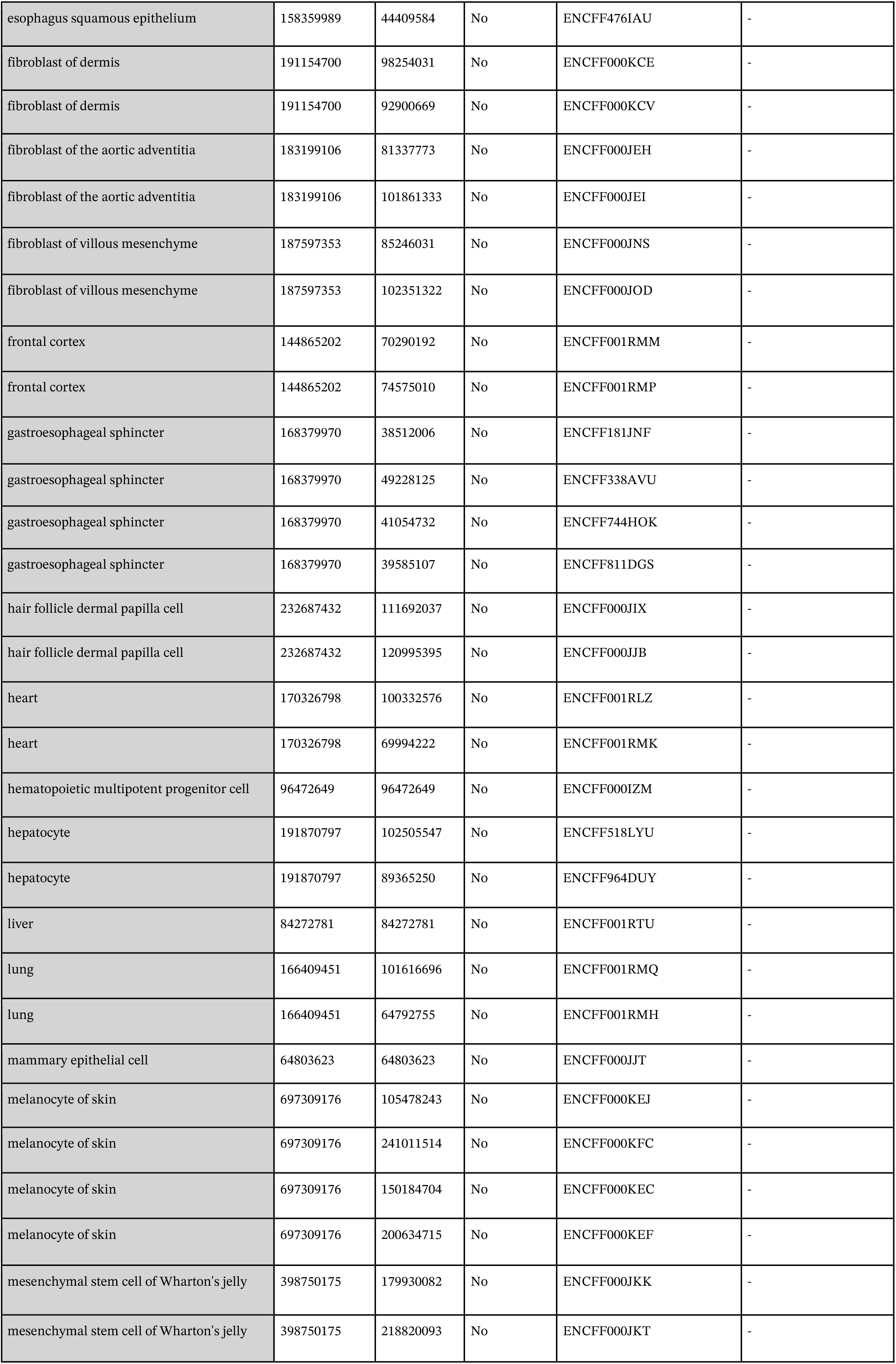
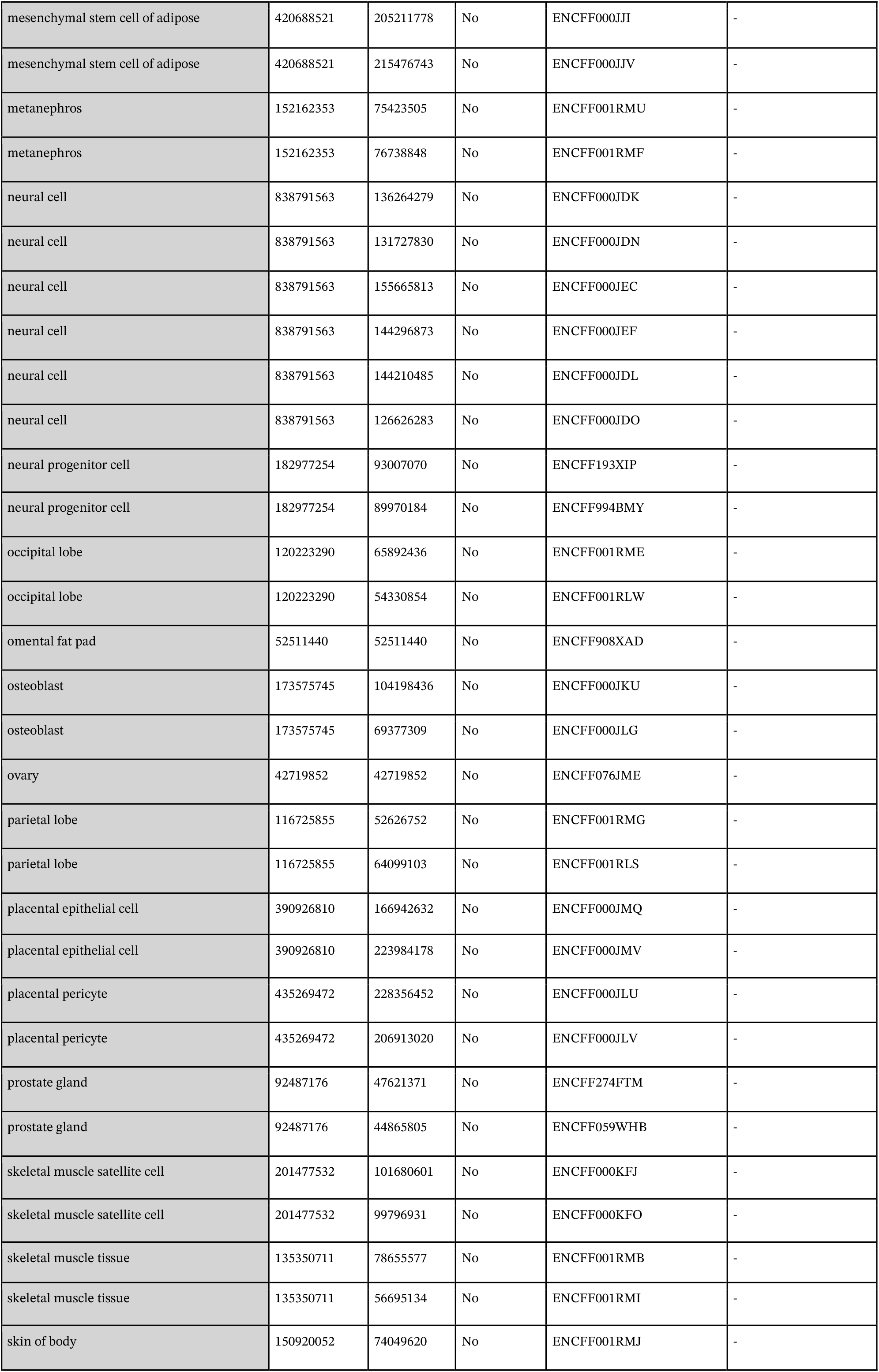
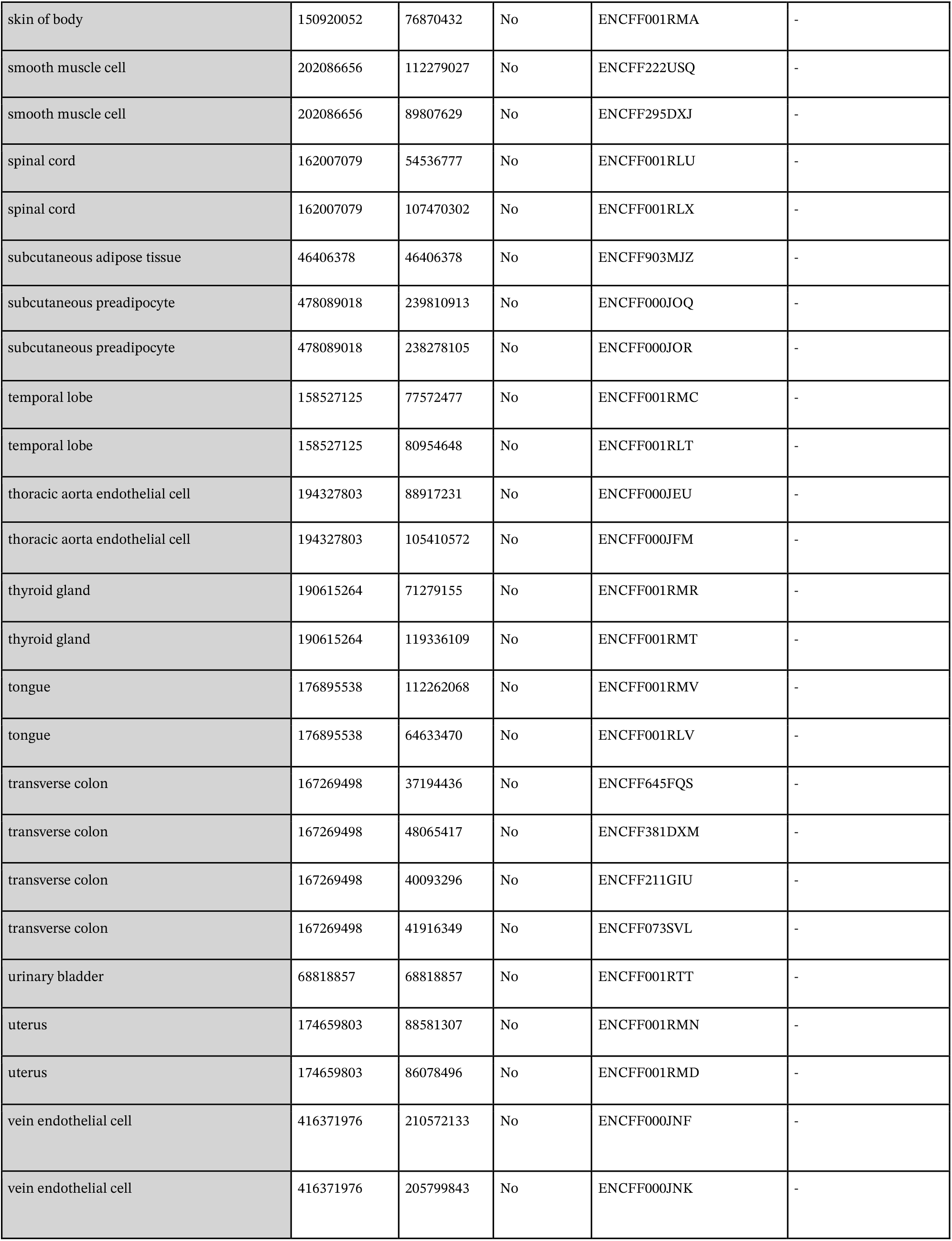
small RNA-seq data used in this study.

